# A conserved MFS orchestrates a subset of O-glycosylation to facilitate macrophage dissemination and tissue invasion

**DOI:** 10.1101/415547

**Authors:** Katarína Valošková, Julia Biebl, Marko Roblek, Shamsi Emtenani, Attila Gyoergy, Michaela Mišová, Aparna Ratheesh, Kateryna Shkarina, Ida S.B. Larsen, Sergey Y. Vakhrushev, Henrik Clausen, Daria E. Siekhaus

## Abstract

Aberrant display of the truncated core1 O-glycan T-antigen is a common feature of human cancer cells that correlates with metastasis. Here we show that T-antigen in *Drosophila melanogaster* macrophages is involved in their developmentally programmed tissue invasion. Higher macrophage T-antigen levels require an atypical major facilitator superfamily (MFS) member that we named Minerva which enables macrophage dissemination and invasion. We characterize for the first time the T and Tn glycoform O-glycoproteome of the *Drosophila melanogaster* embryo, and determine that Minerva increases the presence of T-antigen on protein pathways previously linked to cancer, most strongly on the protein sulfhydryl oxidase Qsox1 which we show is required for macrophage invasion. Minerva’s vertebrate ortholog, MFSD1, rescues the *minerva* mutant’s migration and T-antigen glycosylation defects. We thus identify a key conserved regulator that orchestrates O-glycosylation on a protein subset to activate a program governing migration steps important for both development and cancer metastasis.

## INTRODUCTION

The set of proteins expressed by a cell defines much of its potential capacities. However, a diverse set of modifications can occur after the protein is produced to alter its function and thus determine the cell’s final behavior. One of the most frequent, voluminous and variable of such alterations is glycosylation, in which sugars are added onto the oxygen (O) of a serine or threonine or onto the nitrogen (N) of an asparagine (Kornfeld and Kornfeld, 1985; Marshall, 1972; Ohtsubo and Marth, 2006). O-linked addition can occur on cytoplasmic and nuclear proteins in eukaryotes (Comer and Hart, 2000; Hart et al., 2011), but the most extensive N- and O-linked glycosylation occurs during the transit of a protein through the secretory pathway. A series of sugar molecules are added starting in the endoplasmic reticulum (ER) or cis-Golgi and continuing to be incorporated and removed until passage through the trans Golgi network is complete (Aebi, 2013; Stanley et al., 2009). N-linked glycosylation is initiated in the ER at consensus NxS/T X≠ P site, whereas the most common GalNAc-type O-linked glycosylation is initiated in the early Golgi and glycosites display no clear sequence motifs, apart from a prevalence of neighboring prolines (Bennett et al., 2012; Christlet and Veluraja, 2001). Glycosylation can affect protein folding, stability and localization as well as serve specific roles in fine-tuning protein processing and functions such as protein adhesion and signaling (Goth et al., 2018; Varki, 2017). The basic process by which such glycosylation occurs has been well studied. However our understanding of how specific glycan structures participate in modulating particular cellular functions is still at its beginning.

The need to understand the regulation of O-glycosylation is particularly relevant for cancer (Fu et al., 2016; Häuselmann and Borsig, 2014). The truncated O-glycans called T and Tn antigen are not normally found on most mature human cells (Cao et al., 1996) but up to 95% of cells from many cancer types display these at high levels (Boland et al., 1982; Cao et al., 1996; Howard and Taylor, 1980; Limas and Lange, 1986; Orntoft et al., 1985; Springer, 1984; Springer et al., 1975). The T O-glycan structure (Galβ1-3GalNAcα1-O-Ser/Thr) is synthesized by the large family of polypeptide GalNAc-transferases (GalNAc-Ts) that initiate protein O-glycosylation by adding GalNAc to form Tn antigen and the core1 synthase C1GalT1 that adds Gal to the initial GalNAc residues (Tian and Ten Hagen, 2009) to form T antigen (**Fig 1A**). The human C1GalT1 synthase requires a dedicated chaperone, COSMC, for folding and ER exit (Ju and Cummings, 2005). In adult humans these O-glycans are normally capped by sialic acids and/or elongated and branched into complex structures (Tarp and Clausen, 2008). However, in cancer this elongation and branching is reduced or absent and the appearance of these truncated T and Tn O-glycans correlates positively with cancer aggressiveness and negatively with long-term prognoses for many cancers in patients (Baldus et al., 2000; Carrasco et al., 2013; Ferguson et al., 2014; MacLean and Longenecker, 1991; Schindlbeck et al., 2005; Springer, 1997, 1989; Summers et al., 1983; Yu et al., 2007). The molecular basis for the enhanced appearance of T antigen in cancers is not clear (Chia et al., 2016), although higher Golgi pH in cancer cells correlates with increases in T antigen (Kellokumpu, Sormunen and Kellokumpu, 2002). Interestingly, T antigen is also observed as a transient fetal modification (Barr et al., 1989) and cancer cells frequently recapitulate processes that happened earlier in development (Cofre and Abdelhay, 2017; Pierce, 1974). Identifying new mechanisms that regulate T antigen modifications developmentally has great potential to lead to important insights into cancer biology.

**Figure 1:**
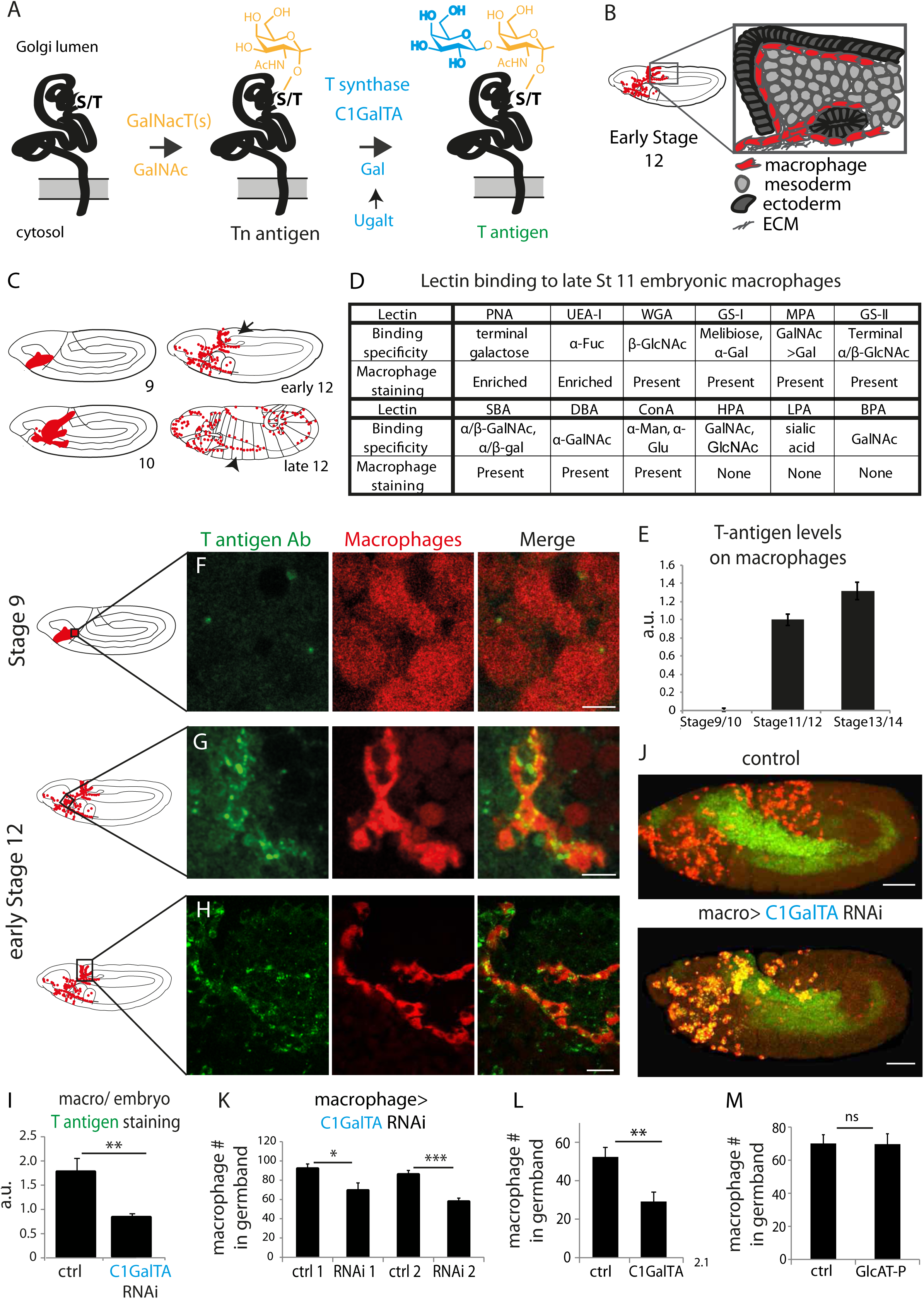
T antigen is enriched on *Drosophila* macrophages prior to and during their invasion of the extended germband. (**A**) Schematic of T antigen modification of serine (S) and threonine (T) on proteins within the Golgi lumen, through successive addition of GalNAc (yellow) by GalNAcTs and Gal (blue) by C1GalTs. Ugalt transports Gal into the Golgi. Glycosylation is shown at a much larger scale than the protein. (**B**) Schematic of an early Stage 12 embryo and a magnification of macrophages (red) entering between the germband ectoderm (dark grey), and mesoderm (light grey). (**C**) Schematic showing macrophages (red) disseminating from the head mesoderm in Stage 9. By Stage 10, they migrate towards the extended germband, the dorsal vessel and along the ventral nerve cord (vnc). At late Stage 11 germband invasion (arrow) begins and continues during germband retraction. Arrowhead highlights migration along the vnc in late Stage 12. **(D)** Table summarizing a screen of glycosylation-binding lectins for staining on macrophages invading the germband in late Stage 11 embryos. Enrichment was seen for PNA which recognizes T antigen and UEA-I which recognizes fucose. (**E**) Quantification of T antigen fluorescence intensities on wild type embryos shows upregulation on macrophages between Stage 9/10 and Stage 11/12. Arbitrary units (au) normalized to 1 for Stage 11) p <0.0001. (**F-H**) Confocal images of fixed lateral wild type embryos from (**F**) Stage 9 and (**G-H**) early Stage 12 with T antigen visualized by antibody staining (green) and macrophages by *srpHemo-3xmCherry* expression (red). Schematics at left with black boxes showing the imaged regions. (**I**) Quantification of control shows T antigen enrichment on macrophages when normalized to whole embryo. RNAi in macrophages against C1GalTA by *srpHemo(macrophage)>C1GalTA RNAi vdrc2826* significantly decreases this T antigen staining (n=8 embryos, p= 0.0107). (**J**) Representative confocal images of Stage 12 embryos from control and the aforementioned C1GalTA RNAi. Macrophages marked with cytoplasmic GFP (red) and nuclear RFP (green). **(K,L)** Quantification of macrophages in the germband in Stage 12 embryos for (**K**) control and two independent RNAis against C1GalTA (*vdrc110406 or vdrc2826*) expressed in macrophages by the *srpHemo-Gal4* driver (n=21-31 embryos,p <0.0001 and 0.0174) or (**L**) in control and the *C1GalTA[2.1]* excision mutant (n=23-24, p=0.0006). Macrophages labeled with *srpHemo-H2A∷3xmCherry.* The RNAis and the mutant significantly decreased the macrophage number, arguing that T antigen is required in macrophages for germband entry. (**M**) Quantification of germband macrophages in early Stage 12 embryos in control and *GlcAT-P*^*MI05251*^ embryos shows no defect in macrophage invasion in the mutant (n=17-20, p=0.9617). **E** analyzed by Kruskal-Wallis test **I, K-M** analyzed by Student’s t-test. ns=p>0.05, * p<0.05; ** p<0.01; *** p<0.001. Scale bars represent 50µm in **J,** and 10µm in **F-H**. See also Fig S1.

*Drosophila* as a classic genetic model system is an excellent organism in which to investigate these questions. *Drosophila* displays T antigen as the predominant form of GalNAc-, or mucin-type, O-glycosylation in the embryo with 18% of the T glycans being further elaborated, predominantly by the addition of GlcA (Aoki et al., 2008). As in vertebrates, the GalNAc-T isoenzymes directing the initial step of GalNAc addition to serines and threonines are numerous, with several already known to display conserved substrate specificity *in vitro* with vertebrates (Müller et al., 2005; Schwientek et al., 2002; Ten Hagen et al., 2003). The *Drosophila* GalNAc-Ts affect extracellular matrix (ECM) secretion, gut acidification and the formation of the respiratory system (Tian and Ten Hagen, 2006; Tran et al., 2012; Zhang et al., 2010). In flies the main enzyme adding Gal to form T antigen is C1GalTA (Müller et al., 2005) whose absence causes defects in ventral nerve cord (vnc) condensation during Stage 17, hematopoetic stem cell maintenance, and neuromuscular junction formation (Fuwa et al., 2015; Itoh et al., 2016; Lin et al., 2008; Yoshida et al., 2008). While orthologous to the vertebrate Core1 synthases, the *Drosophil*a C1GALTs differ in not requiring a specific chaperone (Müller et al., 2005). Most interestingly, the T antigen is found on embryonic macrophages (Yoshida et al., 2008), a cell type which can penetrate into tissues in a manner akin to metastatic cancer (Ratheesh et al., 2018; Siekhaus et al., 2010). Macrophage invasion of the germband (**Fig 1B**, arrow in **Fig 1C**) occurs between the closely apposed ectoderm and mesoderm (Ratheesh et al., 2018; Siekhaus et al., 2010) from late Stage 11 through Stage 12 during the dispersal of macrophages throughout the embryo (**Fig 1C**) along routes that are mostly noninvasive, such as along the inner ventral nerve cord (vnc) (arrowhead in **Fig 1C**) (Campos-Ortega and Hartenstein, 1997; Evans et al., 2010). Given these potentially related but previously unconsolidated observations, we sought to determine the relationship between the appearance of T antigen and macrophage invasion and to use the genetic power of *Drosophila* to find new pathways by which this glycophenotype is regulated.

## RESULTS

### T antigen is enriched and required in invading macrophages in *Drosophila* embryos

To identify glycan structures present on macrophages during invasion we performed a screen examining FITC-labelled lectins (see Methods for abbreviations). Only two lectins had higher staining on macrophages than on surrounding tissues (labeled enriched): PNA, which primarily binds to the core1 T O-glycan, and UEA-I, which recognizes Fucα1-2Galβ1-4GlcNAc(Molin et al., 1986; Natchiar et al., 2007) (**Fig 1D, S1A-B).** Both glycans are associated with the invasive migration of cancer cells (Agrawal et al., 2017; Hung et al., 2014). SBA, WGA, GS-II, GS-I, ConA, MPA and BPA bound at similar or lower levels on macrophages compared to flanking tissues (**Fig 1D, S1C-I**). We saw no staining with the sialic acid-recognizing lectin LPA, and none with DBA and HPA, that both recognize αGalNAc (Piller et al., 1990) (**Fig 1D, S1K-L**). Thus T antigen and a fucosylated structure are upregulated on embryonic macrophages during their invasion. To confirm T antigen as the source of the macrophage signal, and to characterize its temporal and spatial enrichment, we used a monoclonal antibody (mAb 3C9) to the T O-glycan structure (Steentoft et al., 2011). Through Stage 10, macrophages displayed very little T antigen staining, similar to other tissues (**Fig 1E, F**). However, at late Stage 11 (**Fig S1A**) and early Stage 12, when macrophages start to invade the extended germband, T antigen staining began to be enriched on macrophages moving towards and into the germband (**Fig 1E-H**). We knocked down the core1 synthase C1GalTA required for the final step of T antigen synthesis (**Fig 1A**) (Lin et al., 2008; Müller et al., 2005) using RNAi expression only in macrophages and observed strongly reduced staining (**Fig 1I, Fig S1M**). We conclude that the antibody staining is the result of T antigen produced by macrophages themselves. Our results are consistent with findings showing T antigen expression in a macrophage-like pattern in late Stage 12, and on a subset of macrophages at Stage 16 (Yoshida et al., 2008). To determine if these T O-glycans on macrophages are important for facilitating their germband invasion, we knocked down C1GalTA in macrophages with two independent RNAi lines, and used a P element excision allele, C1GalTA[2.1] which removes conserved sequence motifs required for activity (Lin et al., 2008). We visualized macrophages through specific expression of fluorescent markers and observed a 25 and a 33% decrease in their number in the germband for the RNAis (**Fig 1J,K),** and a 44% decrease in the C1GalTA[2.1] mutant (**Fig 1L**). When we counted the number of macrophages sitting on the yolk next to the germband in the strongest RNAi we observed an increase (**Fig S1N**) that we also observed in the C1GalT mutant (**Fig S1O**). The sum of the macrophages in the yolk and germband is the same in the control, RNAi knockdown (control 136.5±6.4, RNAi 142.3±6.6, p=0.7) and mutant (control 138.5±4.9, mutant, 142.3±7.4, p=0.87) arguing that macrophages that cannot enter the germband when C1GalTA levels are reduced remain on the yolk (**Fig S1O)**. We observed no effect on the migration of macrophages on the vnc, a route that does not require tissue invasion (**Fig S1P**) (Campos-Ortega and Hartenstein, 1997; Evans et al., 2010). 18% of T antigen in the embryo has been found to be further modified, predominantly by glucoronic-acid (GlycA) (Aoki et al., 2008). Of the three GlcA transferases found in *Drosophila* only GlcAT-P is robustly capable of adding GlcA onto the T O-glycan structure in cells (Breloy et al., 2016; Itoh et al., 2018; Kim et al., 2003). To examine if the specific defect in germband invasion that we observed by blocking the formation of T antigen is due to the need for a further elaboration by GlcA, we utilized a lethal MI{MIC} transposon insertion mutant in the GlcAT-P gene. We observed no change in the numbers of macrophages within the germband in the GlcAT-PMI05251 mutant (**Fig 1M**) and a 20% increase in the number of macrophages on the yolk (**Fig S1P**). Therefore our results strongly suggest that the T antigen we observe being upregulated in macrophages as they move towards and into the germband is needed for efficient tissue invasion.

### An atypical MFS member acts in macrophages to increase T antigen levels

We sought to determine which proteins could temporally regulate the increase in the appearance of T O-glycans in invading macrophages. We first considered proteins required for synthesizing the core1 structure, namely the T synthase, C1GalTA, and the UDP-Gal sugar transporter, Ugalt (Aumiller and Jarvis, 2002) (**Fig 1A**). However, q-PCR analysis of FACS sorted macrophages from Stage 9-10, Stage 12, and Stage 13-17 show that though both are enriched in macrophages, neither is transcriptionally upregulated before or during Stage 12 (**Fig 2A,B**). We therefore examined the Bloomington *Drosophila* Genome Project (BDGP) *in situ* database looking for predicted sugar binding proteins expressed in macrophages with similar timing to the observed T antigen increase (Tomancak et al., 2007, 2002). We identified CG8602, a predicted MFS with regions of homology to known sugar responsive proteins and predicted sugar or neurotransmitter transporters (**Fig 2C).** BDGP and our *in situ* hybridizations indicate that CG8602 RNA is maternally deposited, with expression throughout the embryo through Stage 4 after which its levels decrease (**Fig S2A**). This weak ubiquitous expression is followed by strong enrichment in macrophages from Stage 10-12 (**Fig 2D**), along with expression in the amnioserosa at Stage 13 (**Fig S2B**). We confirmed this by q-PCR analysis of FACS sorted macrophages, which detected seven-fold higher levels of CG8602 RNA in macrophages than in the rest of the embryo by Stage 9-10 and 12-fold by Stage 12 (**Fig 2E**). To determine if CG8602 could affect T antigen levels, we examined a viable P-element insertion mutant in the 5’UTR, *CG8602*^*EP3102*^ (**Fig S2C**). This insertion displays strongly reduced CG8602 expression in FACS-sorted macrophages to 15% of wild type levels, as assessed by q-PCR (**Fig 2F**), and shows strongly diminished expression throughout the embryo by *in situ* hybridization (**Fig S2D**). We also created an excision allele, Δ33, removing the 5’UTR flanking the P-element, the start methionine, and 914 bp of the ORF (**Fig S2C**). This is a lethal allele, and the line carrying it over a balancer is very weak; exceedingly few embryos are laid and the embryos homozygous for the mutation do not develop past Stage 12. Therefore, we did not continue experiments with this allele, and instead utilized the insertion mutant. This *CG8602*^*EP3102*^ P-element mutant displays decreased T antigen staining on macrophages moving towards and entering the germband (**Fig 2G**) in Stage 11 through late Stage 12. q-PCR analysis on FACS sorted macrophages show that the reduction in T antigen levels in the mutant is not caused by changes in the RNA levels of the T synthase C1GalTA or the Ugalt Gal and GalNAc transporter (Aumiller and Jarvis, 2002; Segawa et al., 2002) (**Fig 2H**). Since O-glycosylation is initiated in the Golgi, we wanted to examine where CG8602 is localized. We first utilized the macrophage-like S2R+ cell line, transfecting a FLAG∷HA or 3xmCherry labeled form of CG8602 under srpHemo or a copper inducible MT promoter control. We detected no colocalization with markers for the nucleus, ER, peroxisomes, mitochondria or lysosomes (**Fig S2E,J-L)**, but did with the Golgi marker Golgin 84 and the endosome markers Rab7, Rab11 and Hrs (Riedel et al., 2016) (**Fig S2F-I**). We confirmed this Golgi and endosome colocalization with Golgin 84 and Hrs in late Stage 11 embryos using macrophages extracted from positions in the head adjacent to the germband (**Fig 2I**). We conclude that the T antigen enrichment on macrophages migrating towards and into the germband requires a previously uncharacterized atypical MFS with homology to sugar binding proteins that is localized predominantly to the Golgi and endosomes.

**Figure 2:**
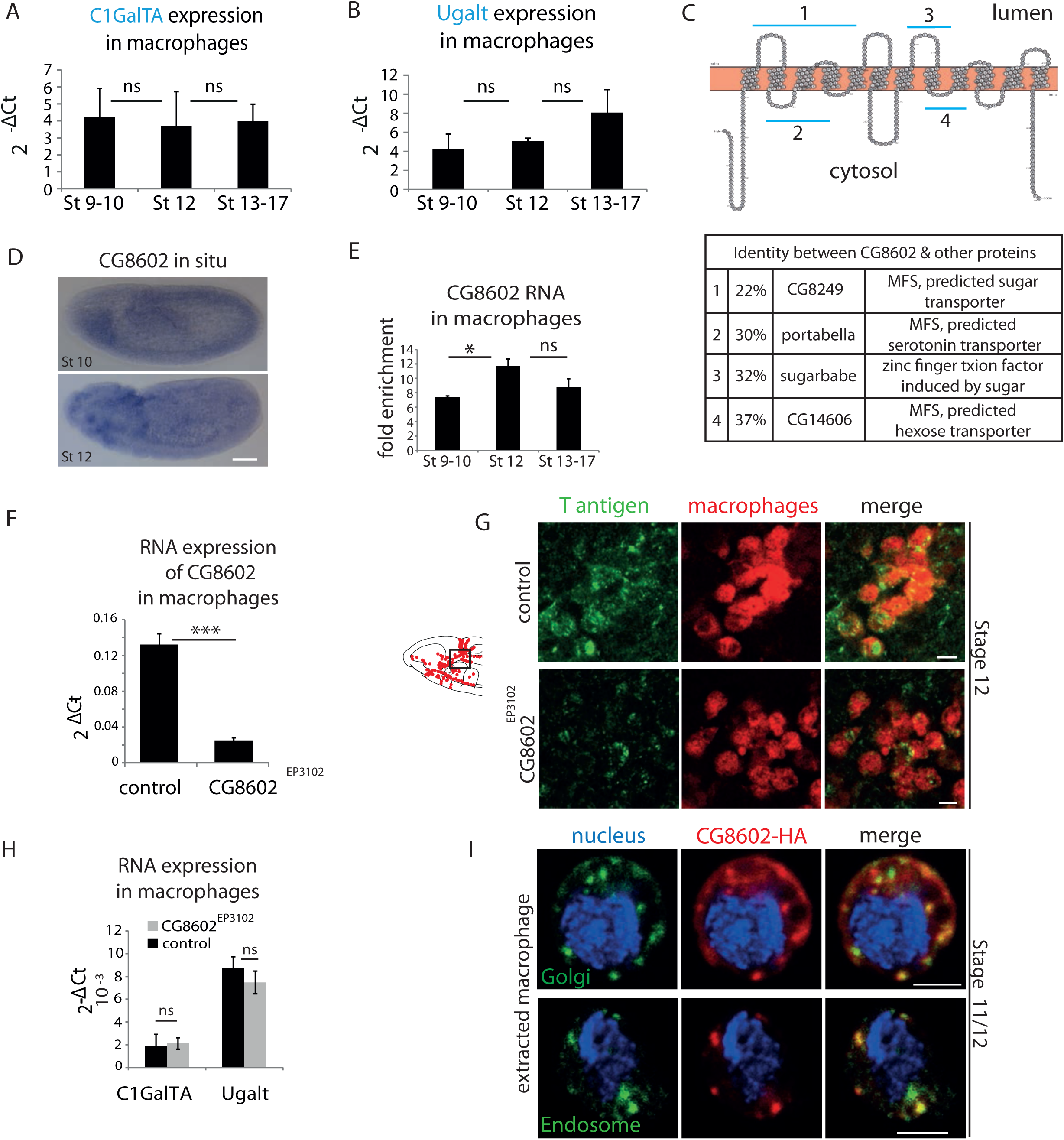
An atypical MFS family member, CG8602, located in the Golgi and endosomes, is required for T antigen enrichment on invading macrophages. (**A,B**) qPCR quantification (2^-ΔCt^) of RNA levels in *mCherry+* macrophages FACS sorted from *srpHemo-3xmCherry* wild type embryos reveals no significant change in the expression of (**A**) the C1GalTA galactose transferase or (**B**) the Ugalt Gal transporter during Stage 9-17 (n=7 biological replicates, 3 independent FACS sorts). **(C)** Schematic made with Protter (Omasits et al., 2014) showing the predicted 12 transmembrane domains of CG8602. Blue lines indicate regions displaying higher than 20% identity to the correspondingly numbered *Drosophila* protein indicated below, along with the homologous protein’s predicted or determined function. **(D**) *In situ* hybridizations of wild type lateral embryos reveal enriched CG8602 expression in macrophages in Stage 10 and 12 and in the amnioserosa by Stage 12 along with lower level ubiquitous expression. (**E**) Quantification by qPCR of CG8602 RNA levels in FACS sorted *mCherry+* macrophages compared to other *mCherry-*cells obtained from *srpHemo-3xmCherry* wild type embryos at Stage 9-10, Stage 12 and Stage 13-17. CG8602 macrophage expression peaks at Stage 12, during macrophage germband entry (n=3-7 biological replicates, 4 independent FACS sorts). **(F)** qPCR quantification in FACS sorted *srpHemo-3xmCherry* labeled macrophages from control and *CG8602*^*EP3102*^ mutant Stage 12 embryos shows an extremely strong decrease in CG8602 RNA expression in the P element insertion mutant used in this study (n=7 biological replicates, 3 independent FACS sorts). **(G**) Confocal images of Stage 12 control and *CG8602*^*EP3102*^ mutant embryos with macrophages (red) visualized by *srpHemo-mCherry* expression and T antigen by antibody staining (green). Schematic at left depicts macrophages (red) entering the germband. Black box indicates the region next to the germband imaged at right. We observe decreased T antigen staining on macrophages in the *CG8602*^*EP3102*^ mutant compared to the control. **(H)** qPCR quantification (2^-ΔCt^) of C1GalTA and Ugalt RNA levels in FACS sorted macrophages from Stage 12 embryos from control and *mrva*^*EP3102*^ mutant embryos shows no significant change in expression of the Gal transferase, or the Gal and GalNAc transporter in the mutant compared to the control (n=7 biological replicates, 3 independent FACS sorts). (**I**) Macrophages near the germband extracted from *srpHemo>CG8602-HA* Stage 11/12 embryos show partial colocalization of the HA antibody labeling CG8602 (red) and a Golgin 84 or Hrs antibody marking the Golgi or endosome respectively (green). Nucleus is stained by DAPI (blue). For all qPCR experiments values are normalized to expression of a housekeeping gene RpL32. Scale bars are 50µm in **D**, 5µm in **G,** 3µm in **I**. Significance was assessed by Kruskal-Wallis test in **A, B**, One way Anova in **E** and Student’s t-test in **F, H**. ns=p>0.05, * p<0.05, *** p<0.001. See also Fig S2.

### The MFS, Minerva, is required in macrophages for dissemination and germband invasion

We examined if CG8602 affects macrophage invasive migration. The *CG8602*^*EP3102*^ mutant displayed a 35% reduction in macrophages within the germband at early Stage 12 compared to the control (**Fig 3A,B,D, Fig S3A**). The same decrease is observed when the mutant is placed over the deficiency Df(3L)BSC117 that removes the gene entirely (**Fig3D**), arguing that *CG8602*^*EP3102*^ is a genetic null for macrophage germband invasion. The P element transposon insertion itself causes the migration defect because its precise excision restored the number of macrophages in the germband to wild type levels (**Fig 3D**). Expression of the CG8602 gene in macrophages can rescue the *CG8602*^*EP3102*^ P element mutant (**Fig 3C,D, Fig S3A**), and RNAi knockdown of CG8602 in macrophages can recapitulate the mutant phenotype (**Fig 3I, Fig S3B**). Our data thus argues that CG8602 is required in macrophages themselves for germband invasion.

**Figure 3:**
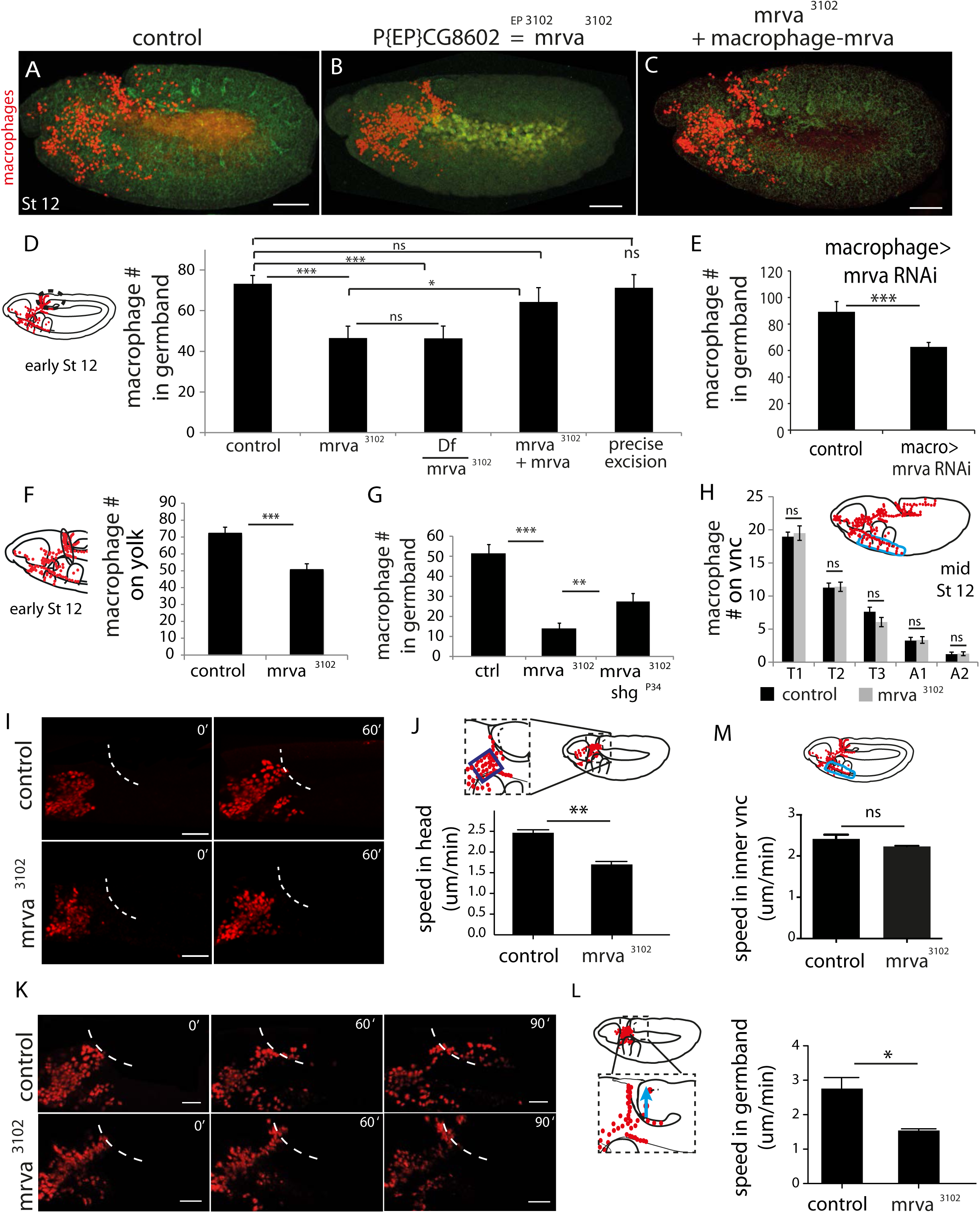
CG8602, which we name Minerva, is required in macrophages for their efficient invasion of the germband. **(A-C)** Representative confocal images of early Stage 12 embryos from **(A**) control, **(B)** *P{EP}CG8602*^*3102*^*=minerva (mrva)*^*3102*^ mutant, and (**C)** *mrva*^*3102*^ mutants with macrophage expression of the gene rescued by *srpHemo(macro)-mrva.* Macrophages express *srpHemo-3XmCherry* (red) and the embryo autofluoresces (green). In the mutant, macrophages remain in the head and fail to enter the germband, hence we name the gene *minerva***. (D)** Dashed ellipse in schematic at left represents the germband region in which macrophage (red) were counted throughout the study. Comparison of the control (n=38), *mrva*^*3102*^ mutants (n=37) and *mrva*^*3102*^ mutant/Df(3L)BSC117 that removes the gene (n=23) shows that the mutant significantly decreases migration into the extended germband. This defect can be partially rescued by expression in macrophages of *srpHemo>mrva∷FLAG∷HA* (n=18) (p<0.05) and completely rescued by precise excision (*mrva*^*Δ32*^) of the P element (n=16). *srpHemo>mcherry-nls* labeled the macrophages. **(E-G)** Macrophage quantification in early Stage 12 embryos. (**E**) Fewer macrophages in the germband are also observed upon expression of mrvaRNAi v101575 only in macrophages under the control of *srpHemo* (n=28-35 embryos). (**F)** Fewer macrophages found on the yolk neighboring the germband (oval in schematic) in the *mrva*^*3102*^ mutant compared to control embryos (n=14-16 embryos, p=0.0003). (**G**) Increased germband macrophage numbers in *shg*^*P*34^, *mrva*^*3102*^ compared to the *mrva*^*3102*^,mutant indicates a partial rescue from reducing DE-Cadherin which is expressed in the germband ectoderm (n=19-29). **(H**) No significant difference in number of macrophages labeled with *srpHemo-3xmcherry* in vnc segments (blue oval in schematic) between control and *mrva*^*3102*^ mutant embryos in fixed mid Stage 12 embryos (n=23-25). Images from two-photon movies of (**I**) Stage 10 and (**K**) late Stage 11-early Stage 12 embryos in which macrophages (red) are labeled with *srpHemo-H2A∷3xmCherry*. (**I**) Stills at 0 and 60 min and (**J**) quantification of macrophage speed reveal 33% slower macrophage migration in the head towards the yolk neighboring the germband in the *mrva*^*3102*^ mutant compared to the control, n=3 movies for each, #tracks: control=329, mutant=340, p=0.002. Blue box in magnification in schematic indicates region analysed in J. (**K**) The time when macrophages reached the germband in each genotype was defined as 0’. Stills at 60 and 90 min and (**L**) quantification of macrophage speed reveal 43% slower macrophage migration in the germband in the *mrva*^*3102*^ mutant compared to the control. Blue arrow in schematic indicates route analyzed. n=3 movies for each, #tracks: control=21, mutant=14, p=0.022. **(M**) Macrophage speed in the inner vnc in early Stage 12 embryos (see schematic above) shows no significant change in the *mrva*^*3102*^ compared to the control, n=3 movies for each, #tracks: control=180, mutant=180, p=0.113. Significance was assessed by Kruskal-Wallis test in **D, G**, Student’s t-test in **E, F, H, J, L,M**. ns=p> 0.05, * p<0.05, ** p<0.01, *** p<0.001. Scale bars are 50µm in A-C, 40µm in **I**, 30µm in **K**. See also Fig S3.

Decreased numbers of macrophages in the extended germband could be caused by specific problems entering this region, or by general migratory defects or a decreased total number of macrophages. To examine the migratory step that precedes germband entry, we counted the number of macrophages sitting on the yolk next to the germband in fixed embryos in the *CG8602*^*EP3102*^ mutant. We observed a 30% decrease compared to the control (**Fig 3F**), suggesting a defect in early dissemination. Entry into the germband by macrophages occurs between the closely apposed DE-Cadherin expressing ectoderm and the mesoderm and is accompanied by deformation of the ectodermal cells(Ratheesh et al., 2018). We tested if reductions in DE-Cadherin could ameliorate the germband phenotype. Indeed, combining the *CG8602*^*EP3102*^ mutation with *shg*^*P34*^ which reduces DE-Cadherin expression (Pacquelet and Røth, 1999; Tepass et al., 1996) produced a partial rescue (**Fig 3G**), consistent with CG8602 playing a role in germband entry as well as an earlier migratory step. Macrophage migration along the vnc in late Stage 12 showed no significant difference in the number of macrophages compared to the control in fixed embryos (**Fig 3H**) from the *CG8602*^*EP3102*^ mutant or from a knockdown in macrophages of *CG8602* by RNAi (**Fig S3C**), arguing against a general migratory defect. There was also no significant difference in the total number of macrophages in either case (**Fig S3D, E).** From analyzing the *CG8602* mutant phenotype in fixed embryos we conclude that CG8602 does not affect later vnc migration but is important for the early steps of dissemination and germband invasion.

To examine the effect of CG8602 on macrophage speed and dynamics, we performed live imaging of macrophages labeled with the nuclear marker *srpHemo-H2A∷3xmCherry* in control and *CG8602*^*EP3102*^ mutant embryos (**Video 1 and 2**). We first imaged macrophages migrating from their initial position in the delaminated mesoderm up to the germband and detected a 33% decrease in speed (2.46±0.07 µm/min in the control, 1.66±0.08 µm/min in the *mrva*^*3102*^ mutant, p=0.002) (**Fig 3I, J**) and no significant decrease in persistence (0.43±0.02 in the control, 0.40±0.01 in the mutant, p=0.218) (**Fig S3F**). We then examined the initial migration of macrophages into the germband at late Stage 11. We observed a range of phenotypes in the six movies we made of the mutant: in half of them macrophages entered at the normal time, and in the other half we observed a one to three hour delay in entry. As we observed no change in the timing of the initiation of germband retraction (269.6±9 min in control and 267.1±3 min in *mutant*, p=-0.75) but did observe a decreased speed of its completion in the mutant (107±12 min from start to end of retraction in control and 133±6 min for mutant p=0.05), we only analyzed macrophages within the germband before its retraction begins. We observed a 43% reduction in macrophage speed within the germband (2.72±0.32 µm/min in the control and 1.55±0.04 µm/min in the mutant, p=0.02) (**Fig 3K,L**). To assess this phenotype’s specificity for invasion, we used live imaging of macrophage migration along the inner vnc that occurs during the same time period as germband entry; we observed no significant change in speed (2.41±0.06 µm/min in the control and 2.23±0.01 µm/min in the mutant, p=0.11) or directionality (0.43±0.03 in the control and 0.43±0.02 in the mutant, p=0.9742) (**Fig 3M**, **Video 3 and 4**). We conclude from the sum of our experiments in fixed and live embryos that CG8602 is important for the initial disseminatory migration out of the head and for invasive migration into and within the germband, but does not alter general migration. We name the gene *minerva (mrva)*, for the Roman goddess who was initially trapped in the head of her father, Jupiter, after he swallowed her pregnant mother who had turned herself into a fly.

### Minerva affects a small fraction of the *Drosophila* embryonic O-glycoproteome

We set out to determine if Minerva induces T glycoforms on particular proteins. We first conducted a Western Blot with a mAb to T antigen on whole embryo extracts. We used the whole embryo because we were unable to obtain enough protein from FACSed macrophages or to isolate CRISPR-induced full knockouts of *minerva* in the S2R+ macrophage-like cell line. We observed that several bands detected with the anti-T mAb were absent or reduced in the *minerva* mutant (**Fig 4A**), indicating an effect on a subset of proteins. We wished to obtain a more comprehensive view of the proteins affected by Minerva. Since there is little information about *Drosophila* O-glycoproteins and O-glycosites (Schwientek *et al.*, 2007; Aoki and Tiemeyer, 2010), we used lectin-enriched O-glycoproteomics to identify proteins displaying T and Tn glycoforms in Stage 11/12 embryos from wild type and *mrva*^*3102*^ mutants (**Fig S4A)**. We labeled tryptic digests of embryonic protein extracts from control or mutant embryos with stable dimethyl groups carrying medium (C_0_H_2_D_4_) or light (C_2_H_6_) isotopes respectively to allow each genotype to be identified in mixed samples(Boersema et al., 2009; Schjoldager et al., 2012, 2015). The pooled extracts were passed over a Jacalin column to enrich for T and Tn O-glycopeptides; the eluate was analyzed by mass spectrometry to identify and quantify T and Tn modified glycopeptides in the wild type and the mutant sample through a comparison of the ratio of the light and medium isotope labeling channels for each glycopeptide. In the wild type we identified T and Tn glycopeptides at 936 glycosites derived from 270 proteins (**Table S1** and **Fig 4B**). 62% of the identified O-glycoproteins and 77% of identified glycosites contained only Tn O-glycans. 33% of the identified O-glycoproteins and 23% of glycosites displayed a mixture of T or Tn O-glycans, and 5% of identified O-glycoproteins and 4% of glycosites had solely T O-glycans (**Fig 4C**). In agreement with previous studies (Steentoft et al., 2013), only one glycosite was found in most of the identified O-glycoproteins (44%) (**Fig 4D**). In 20% we found two sites, and some glycoproteins had up to 27 glycosites. The identified O-glycosites were mainly on threonine residues, (78.5%) with some on serines (21.2%) and very few on tyrosines (0.3%) (**Fig S4B**). Metabolism, cuticle development, and receptors were the most common functional assignments for the glycoproteins (**Fig S4C**). To assess the changes in glycosylation in the *mrva* mutant we utilized two cutoffs, a three-fold and a more stringent ten-fold cutoff. The majority of the quantifiable Tn and T O-glycoproteome was unaltered between the wild type and the *mrva*^*3102*^ mutant, with only 63 proteins (23%) showing more than a three-fold change and 18 (6%) a ten-fold shift (**Fig 4F**). We observed both increases and decreases in the levels of T and Tn modification on proteins in the mutant (**Fig 4F,G, Table S2**), but a greater number of proteins showed decreased than increased T antigen levels. 67% of the vertebrate orthologs of *Drosophila* proteins displaying shifts in this O-glycosylation have previously been linked to cancer (**Fig 4H, Table S2**). These proteins were affected at specific sites, with 40% of glycosites on these proteins changed more than three fold and only 14% more than ten fold. The glycosite shifts in T antigen occurred either without significant alterations in Tn (33% of glycosites had only decreased T antigen, 17% of glycosites had only increased T antigen) or with changes in T antigen occurring in the same direction as the changes in Tn (22% of glycosites both Tn and T antigen increased, 22% of glycosites both Tn and T decreased) (**Table S2**). Only 1% of glycosites displayed decreased T antigen with a significant increase in Tn. Interestingly, a higher proportion of the glycoproteins with altered O-glycosylation in the *mrva*^*3102*^ mutant had multiple glycosites than the general glycoproteome (**Fig 4D**) (P value=0.005 for ten-fold changes). We conclude that Minerva affects O-glycosylation occupancy on a small subset of O-glycoproteins, many of whose vertebrate orthologs have been linked to cancer, with both T and Tn O-glycopeptides being affected.

**Figure 4:**
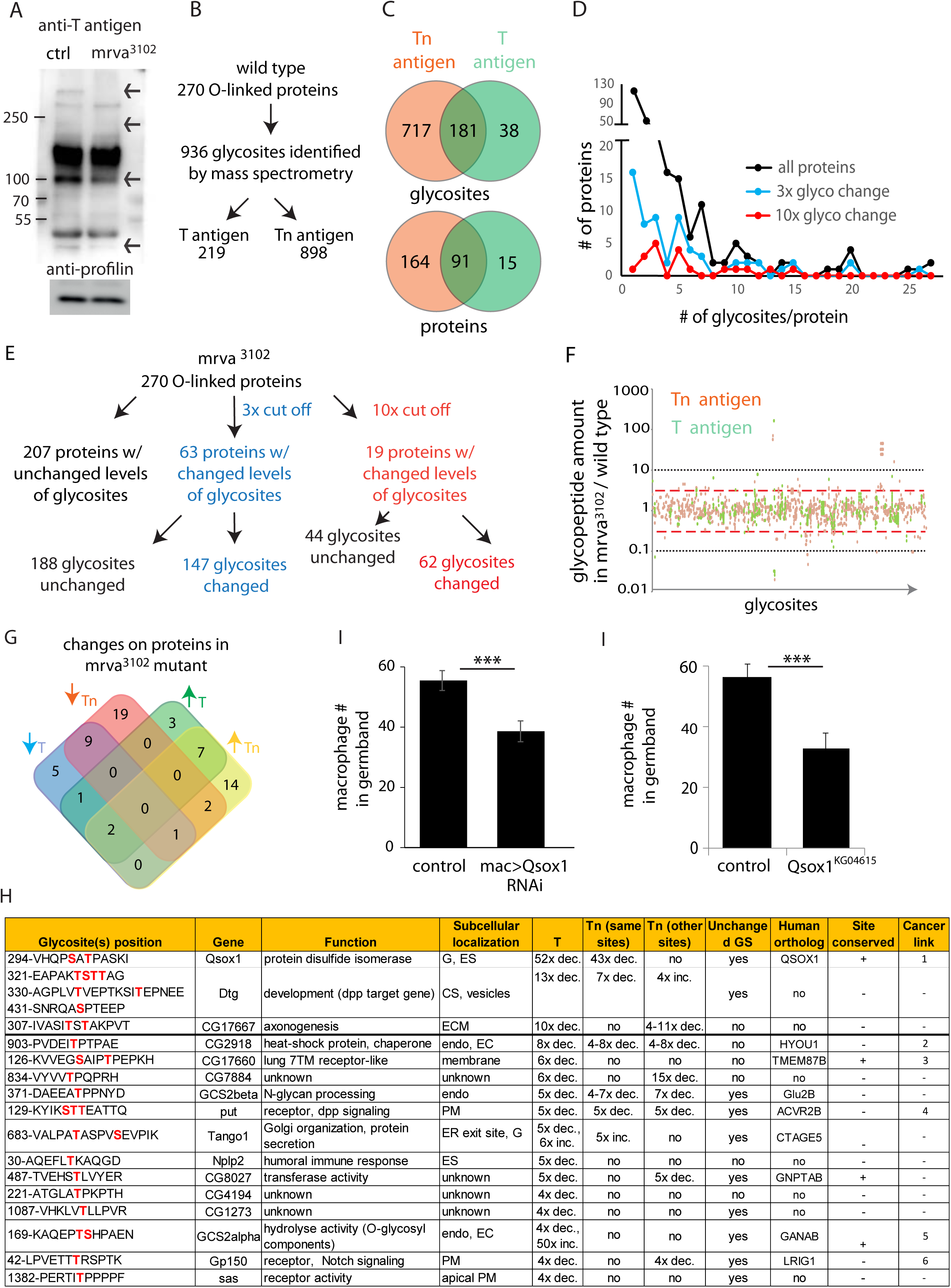
Glycoproteomic analysis reveals Minerva is required for higher levels of T-antigen on a subset of proteins. **(A)** Representative Western blot of protein extracts from Stage 11/12 control and *mrva*^*3102*^ mutant embryos probed with T antigen antibody. Arrows indicate decreased/missing bands in the mutant compared to the control. Profilin serves as a loading control (n=10 biological replicates). **(B)** Summary of glycomics results on wild type embryos. **(C)** Venn diagram indicating number of glycosites or proteins found with T, Tn or T and Tn antigen modifications in the wild type**. (D)** Plot showing the number of T and Tn antigen glycosites per protein in the total glycoproteome and on proteins that show three and ten-fold altered glycopeptides in the *mrva*^*3102*^ mutant. Proteins strongly affected by Minerva have a higher number of glycosites (p = 0.005). (**E**) Summary of glycomics on *mrva*^*3102*^ embryos showing the numbers of proteins and glycosites exhibiting three (blue) or ten (red) fold changes in T and Tn antigen levels. (**F**) T antigen (in orange) and Tn antigen (green) occupied glycosites plotted against the ratio of the levels of glycopeptides found for each glycosite in *mrva*^*3102*^/control mutant. Higher positions on the plot indicate a lower level of glycosylation in the mutant. Red dashed line represents the cut off for 3x changes in glycosylation, and the black dotted line the 10x one. (**G**) Venn diagram of the number of proteins with at least 3 fold change in the T antigen (T) or Tn antigen (Tn) glycosylation in the *mrva*^*3102*^ mutant. Up arrows denote increase, down arrows indicate decrease in levels. (**H**) Proteins with at least a three fold decrease in T antigen levels in the *mrva*^*3102*^ mutant. Glycan modified amino acids are highlighted in bold red font. Unchanged/Higher GS column indicates if any other glycosite on the protein is unchanged or increased. Table does not show the two chitin and chorion related genes unlikely to function in macrophages. G: Golgi, ES: Extracellular space, Endo: Endosomes, ER: Endoplasmic reticulum, ECM: Extracellular Matrix, PM: Plasma Membrane, GS: Glycosite. Cancer links as follows. 1) QSOX1: Promotes cancer invasion *in vitro*, overexpression worse patient outcomes, (Katchman et al., 2013, 2011). 2) HYOU1: Overexpression associated with vascular invasion, worse patient outcomes (Stojadinovic et al., 2007) (Zhou et al., 2016). 3) TMEM87B: translocation breakpoint in cancer, (Hu et al., 2018). 4) ACVR2B: over expressed in renal cancer (Senanayake et al., 2012). 5) GANAB: inhibits cancer invasion *in vitro* (C. Chiu et al., 2011). 6) LRIG1: inhibits cancer invasion *in vitro*, and in mice (Sheu et al., 2014), (Mao et al., 2018). (**I,J**) Quantification in early Stage 12 embryos showing a significant reduction in germband macrophages (**I**) upon the expression in macrophages under *srpHemo-GAL4* of a RNAi line (v108288) against Qsox1 (n=24, 23 embryos) and (**J**) in the P-element mutant KG04615 located in the *Qsox1* 5’UTR. ***, p=0.0006 via Student’s t-test. See also Fig S4, Table S1 and Table S2.

### Minerva raises T antigen levels on proteins required for invasion

Given that the knockdown of the C1GalTA enzyme which blocks Tn to T conversion produced a germband invasion defect, we examined the known functions of the 18 proteins with lower T antigen in the absence of Minerva to distinguish which processes Minerva could influence to facilitate invasion (**Fig 4H**). We excluded two proteins involved in eggshell and cuticle production. To spot proteins whose reduced T antigen-containing glycopeptides are caused directly by alterations in glycosylation rather than indirectly by decreased protein expression in the *mrva* mutant, we checked if glycosylation at other identified glycosites was unchanged or increased. We identified ten such proteins, several of which were in pathways that had been previously linked to invasion in vertebrates. Qsox1, a predicted sulfhydryl oxidase required for the secretion, and thus potential folding of EGF repeats (Tien et al., 2008) showed the strongest alterations of any protein, with a 50-fold decrease in T antigen levels in the *mrva* mutant. The mammalian ortholog has been shown to affect disulfide bond formation, is overexpressed in some cancers, promotes Matrigel invasion, and can serve as a negative prognostic indicator in human cancer patients (Chakravarthi et al., 2007; Katchman et al., 2011; Lake and Faigel, 2014). Dtg, with a 13-fold reduction in T antigen (Hodar et al., 2014), and Put with a five-fold reduction (Letsou et al., 1995) respond to signaling by the BMP-like ligand, Dpp. Gp150 shows a four fold decrease in T antigen and modulates Notch signaling (Fetchko et al., 2002; Li, 2003). Notch and BMP promote invasion and metastasis in mice (Bach et al., 2018; Garcia and Kandel, 2012; Owens et al., 2015; Pickup et al., 2015; Sahlgren et al., 2008; Sonoshita et al., 2011). Dpp signaling directs histoblast invasion in the fly (Ninov et al., 2010). To test if Qsox1, the protein with the strongest changes in T antigen in the *minerva* mutant is required for germband invasion, we examined RNAi knockdown of Qsox1 in macrophages and a P element mutant in the 5’UTR of the Qsox1 gene. In both cases we observed reduced numbers of macrophages in the germband (**Fig 4I,J**) (30% for RNAi and 42% for mutant) and a concomitant increase of macrophages on the neighboring yolk (**Fig S4D,E**). There was no change in total cell number in RNAi knockdown embryos (**Fig S4F**). For technical reasons we did not examine this in the P element mutant line which only grew robustly when combined with a cytoplasmic macrophage marker. We conclude that Mrva is required to increase T O-glycans on a subset of the glycosites of selected glycoproteins involved in protein folding, glycosylation and signaling in pathways frequently linked to promoting cancer metastasis. Its strongest effect is on a predicted sulfhydryl oxidase which is required in macrophages for their germband invasion, the *Drosophila* ortholog of the mammalian cancer protein, QSOX1.

### Conservation of Minerva’s function in macrophage invasion and T antigen modification by its mammalian ortholog MFSD1

To determine if our studies could ultimately be relevant for mammalian biology and therefore also cancer research, we searched for a mammalian ortholog. MFSD1 from *mus musculus*, shows strong sequence similarity with Mrva, with 50% of amino acids displaying identity and 68% conservation (**Fig 5A, Fig S5A**). A transfected C-terminally GFP-tagged form (**Fig S5B**) showed localization to the secretory pathway, colocalizing with the Golgi marker GRASP65 in murine MC-38 colon carcinoma cells (**Fig 5B, Fig S5C-D).** mmMFSD1 expression in macrophages in *mrva*^*3102*^ mutant embryos can completely rescue the germband invasion defect (**Fig 5C,D**). This macrophage-specific expression of MFSD1 also resulted in higher levels of T antigen on macrophages when compared to those in *mrva*^*3102*^ mutants (**Fig 5E,F**). Thus MFSD1 displays localization in the Golgi in mammalian cancer cells and can rescue O-glycosylation and migration defects when expressed in *Drosophila*, arguing that the functions Mrva carries out to promote invasion into the germband are conserved up to mammals.

**Figure 5:**
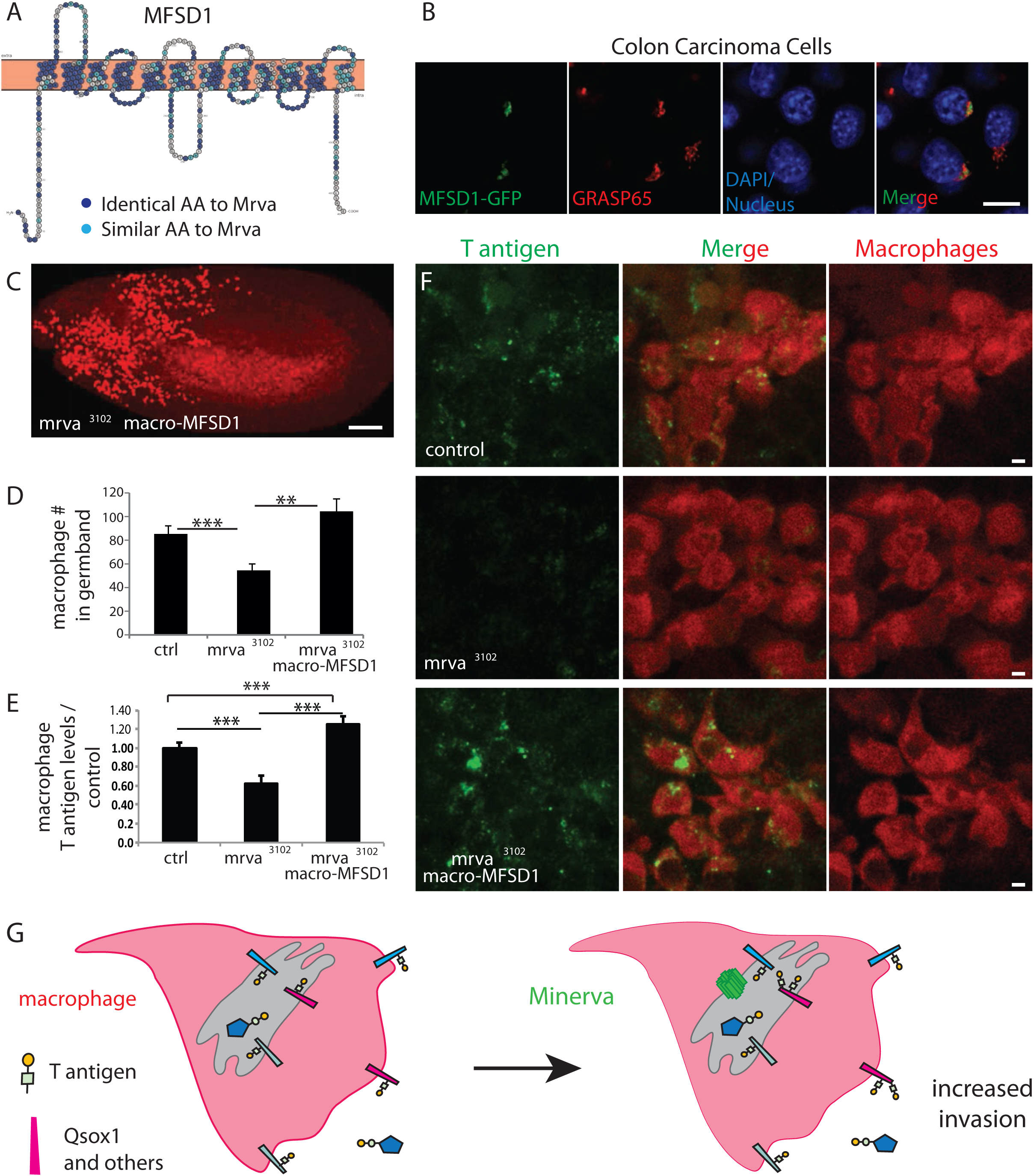
Minerva’s murine ortholog, MFSD1, can substitute for Minerva’s functions in migration and T-antigen glycosylation. (**A**) Topology prediction of mouse MFSD1 (NP_080089.1) using the online tools TMPred (Hofman and Stoffel, 1993) and Protter (Omasits et al., 2014). 50% of amino acids are identical between the *M. musculus* MFSD1 and *D. melanogaster* sequence of *mrva (CG8602)* (NP648103.1) and are highlighted in dark blue, similar amino acids are in light blue. (**B**) Confocal images of MC38 colon carcinoma cells showing colocalization of MFSD1-eGFP (green) with Golgi marker GRASP65 (red). DAPI labels the nucleus (blue). (**C**) Confocal image of a Stage 12 fixed embryo showing that expression of *mmMFSD1* in macrophages under the direct control of the *srpHemo(macro)* promoter in the *mrva*^*3102*^ mutant can rescue the defect in macrophage migration into the germband. Compare to Fig 3A,B. Macrophages visualized with *srpHemo-H2A∷3xmcherry* for C-D. **(D**) Quantitation of the number of macrophages in the germband of early Stage 12 embryos from the control (n=25), *mrva*^*3102*^ mutants (n=29), and *mrva*^*3102*^ *srpHemo(macro)-mmMFSD1* (n=13, p<0.001). (**E**) Quantification of T antigen levels on macrophages in late Stage 11 embryos from control, *mrva*^*3102*^mutant and *mrva*^*3102*^ s*rpHemo(macro)-mmMFSD1* embryos. T antigen levels normalized to those observed in the control (n=8-9 embryos, 280, 333, and 289 cells quantified respectively, p <0.001). **(F)** Confocal images of macrophages (red) on the germband border stained with T antigen antibody (green) in the control, the *mrva*^*3102*^ mutant, and *mrva*^*3102*^ s*rpHemo(macro)-mmMFSD1* shows that mmMFSD1 expression in macrophages can rescue the decrease of macrophage T antigen observed in the *mrva*^*3102*^ mutant. Macrophages visualized with *srpHemo-3xmcherry* for E-F. (**G**) Model for Minerva’s function during macrophage invasion. Minerva in the Golgi (grey) leads to increases in T antigen levels on a subset of proteins that aid invasion, including Qsox1, a sulfhydryl oxidase that regulates protein folding through disulfide bond isomerization. Significance was assessed by Kruskal-Wallis test in **D,E**. ***=p<0.001. Scale bars are 10µm in B, 50µm in D, and 3µm in F. See also Fig S5.

## Discussion

O-glycosylation is one of the most common posttranslational modifications, yet the intrinsic technical challenges involved in identifying O-glycosites and altered O-glycosylation on a proteome-wide level has hampered the discovery of biological functions (Levery et al., 2015). Here we provide two important new advances for the field: (i) defining the GalNAc-type O-glycoproteome of *Drosophila* embryos and (ii) identifying a key regulator of this O-glycosylation, Minerva, with an unexpected role for a member of the major facilitator superfamily. As O-glycosites cannot as yet be reliably predicted, our proteomic characterization in a highly genetically accessible organism will permit future studies on how glycosylation affects cell behavior; we highlight T and Tn O-glycosylated receptors in **Table 1** to further this goal. Our demonstration that a conserved protein affects invasion and the appearance of the cancer-associated core1 T glycoform on a set of proteins connected to invasion may have implications for cancer.

**Table 1:** Receptors identified by the O-glycoproteome as T or Tn antigen modified.

| Receptor | Function | Glycosylation | Changes in *mrva*<sup>3102</sup> |
| --- | --- | --- | --- |
| Babo | Activating signaling | 2 glyco sites, T antigen only | no |
| Boi | Regulation of hh-dependent processes | 3 glyco sites, Tn antigen only | no |
| CG12121 | Unknown | 3 glyco sites, Tn antigen only | no |
| CG15765 | Carbohydrate binding, nervous system development | 1 glycosite, T antigen | no |
| CG5888 | Unknown | 1 glyco site, T or Tn antigen | no |
| CG9095 | Carbohydrate binding | 1 glyco site, Tn antigen | no |
| Cirl | Calcium independent receptor for a-latrotoxin, adult locomotory behavior | 1 glyco site, T or Tn antigen | no |
| Crb | epithelial morphogenesis, apico-basal cell polarity, negative regulator of Notch activity | 1 glyco site, Tn antigen | no |
| Dg | non-integrin ECM receptor, connects ECM to the actin cytoskeleton | 1 glycosite, T or Tn antigen | no |
| Drl | axon guidance through Wnt5 | 1 glyco site, T or Tn antigen | no |
| Hbs | Muscle cell fusion | 2 glycosites, T and Tn antigen | no |
| Hmu | Hydrolase activity | 15 glycosites, both T and Tn antigen | Tn inc. |
| LpR1 | Regulation of immune responses | 2 glycosites, Tn antigen | Tn inc. |
| LpR2 | Cellular uptake of neutral lipids | 3 sites, T and Tn antigen | T & Tn inc. |
| LRP1 | LDL receptor, works with megalin | 4 glycosites, T and Tn antigen | no |
| Mgl | Lipid regulation | 2 glycosites, Tn antigen | Tn dec. |
| Mthl5 | GPCR, heart morphogenesis | 1 glyco site, T or Tn antigen | no |
| NimB2 | Defense response to bacterium | 1 glycosite, Tn antigen | no |
| NimC4 | Recognition and engulfment of apoptotic cells during development | 1 glycosite, T or Tn antigen | no |
| Nrx-IV | Septate junction formation, glial neural interaction | 1 glyco site, Tn antigen | no |
| PlexB | Axon guidance | 1 glyco site, Tn antigen | no |
| Put | Dpp signaling | 5 glyco sites, T and Tn antigen | T&Tn dec. |
| Sas | Pathfinding, glial neuron interaction |  | T dec. |
| Sdc | Robo neural pathfinding, synapse at neuromascular junction | 1 glyco site, Tn antigen | no |
| Sema-1b | Neural pathfinding | 1 glyco site, Tn antigen | no |
| Sli | Neural pathfinding, robo interaction | 2 glyco sites, T and Tn antigen | T&Tn inc. |
| Sr-CII | Scavenger receptor, immune response | 6 glyco sites, T and Tn antigen | no |
| Syb | Synaptic vesicle, SNAP receptor activity | 1 glyco site, T or Tn antigen | no |
| Tequila | Scavenger receptor, serine protease, glucose homeostasis, long and short term memory | 5 glycosite, Tn antigen | no |
| Unc-5 | Neural pathfinding, netrin receptor | 1 glycosite, Tn antigen | no |
| Verm | Cuticle development and tracheal tube size control | 1 glycosite, T or Tn antigen | T &Tn inc. |

### Modifications of the O-glycoproteome by an MFS family member

Our identification of a MFS family member as a regulator of O-glycosylation is surprising. MFS family members can serve as transporters and shuttle a wide variety of substrates (Quistgaard et al., 2016; Reddy et al., 2012). Minerva is localized to the Golgi and displays homology to sugar transporters; Minerva could thus affect O-glycosylation through substrate availability. However, the lower and higher levels of glycosylation in the *mrva*^*3102*^ mutant we observe are hard to reconcile with this hypothesis. Given that the changes in T antigen on individual glycosites in the *mrva* mutant are found either with no significant change in Tn or with a change in the same direction (**Table S2**), regulation appears to occur at the initial GalNAc addition on the protein subset as well as on further T antigen elaboration. 95% of the proteins with 10-fold altered glycosylation in the *mrva* mutant had multiple O-glycosylation sugar modifications compared to 56% of the general O-glycoproteome. Greatly enhanced glycosylation of protein sequences containing an existing glycan modification is observed for some GalNAc-Ts due to a lectin domain(Hassan et al., 2000; Kubota et al., 2006; Revoredo et al., 2016) and Minerva could affect such a GalNAc-T in *Drosophila*. Alternatively, Minerva, while in the “outward open” conformation identified for MFS structures (Quistgaard et al., 2016), may itself have a lectin-like interaction with Tn and T glycoforms that have already been added on a loop of particular proteins. Minerva’s binding could open up the target protein’s conformation to increase or block access to other potential glycosites and thus affect the final glycosylation state on select glycoproteins.

The changes we see in O-glycosylation are also likely due to a combination of Minerva’s direct and indirect effects. O-GalNAc modification of vertebrate Notch can affect Notch signaling during development (Boskovski et al., 2015); the *Drosophila* ortholog of the responsible GalNAc transferase is also essential for embryogenesis (Bennett et al., 2010; Schwientek et al., 2002). Thus the changed glycosylation we observe on components of the Notch and Dpp pathways could alter transcription (Hamaratoglu et al., 2014; Ntziachristos et al., 2014), shifting protein levels and thereby changing the ratio of some glycopeptides in the *mrva* mutant relative to the wild type. Proteins in which glycosylation at other sites is unchanged or changed in the opposite direction are those most likely to be directly affected by Minerva. Such proteins include ones involved in protein folding and O-glycan addition and removal (**Fig 4I**) (Tien et al., 2008). If changes in the glycosylation of these proteins alters their specificity or activity, some of the shifts we observe in our glycoproteomic analysis could be indirect in a different way; an initial effect of Minerva on the glycosylation of regulators of protein folding and glycosylation could change how these primary Minerva targets affect the glycosylation of a second wave of proteins.

### An invasion program regulated by Minerva

The truncated immature core1 T and Tn O-glycans are not usually present in normal human tissues but exposure of these uncapped glycans has been found on the majority of cancers and serves as a negative indicator of patient outcome (Fu et al., 2016; Springer, 1984). An antibody against T antigen has decreased the metastatic spread of cancer cells in mice (Heimburg et al., 2006). Here we further strengthen the case for a causative relationship between this glycosylation modification and the invasive migration that underlies metastasis. The transient appearance of T antigen in human fetuses (Barr et al., 1989) and the conserved function of Minerva lead us to propose that the change in O-glycosylation in cancer represents the reactivation of an ancient developmental program for invasion. Our embryonic glycoproteome analysis identifies 106 T antigen modified proteins, a very large set to investigate. However, the absence of Mrva causes invasion defects and deficits in T antigen modification on only 10-20 proteins; these include components involved in protein folding, glycosylation modification, and the signaling pathways triggered by Notch and the BMP family member, Dpp. Our working model is that the defect in germband tissue invasion seen in the *mrva* mutant is caused by the absence of T antigen on this group of proteins that act coordinately (**Fig 5G**). 56% of these have vertebrate orthologs, and 55% of those have already been linked to cancer and metastasis. For example, the vertebrate ortholog of Qsox1, the protein with the largest changes in T antigen in the *mrva* mutant which is itself required for germband invasion, enhances cancer cell invasion in *in vitro* assays and higher levels of the protein predict poor patient outcomes (Katchman et al., 2013, 2011). Minerva’s vertebrate ortholog, MFSD1, can rescue macrophage migration defects and restores higher T antigen levels. Tagged versions of Minerva’s vertebrate ortholog, MFSD1, detected the protein in lysosomes in HeLa and rat liver cells (Chapel et al., 2013; Palmieri et al., 2011).

However in cancer cells, we find MFSD1 in the Golgi, where O-glycosylation is known to occur (Bennett et al., 2012). As kinases add phospho-groups to affect a set of proteins and orchestrate a particular cellular response, we propose that Minerva in *Drosophila* macrophages and its vertebrate ortholog MFSD1 in cancer trigger changes in O-glycosylation that coordinately modulate, activate and inhibit a protein group to facilitate cellular dissemination and tissue invasion.

## ACKNOWLEDGEMENTS

We thank the following for their contributions: Dr. McNew and the *Drosophila* Genomics Resource Center supported by NIH grant 2P40OD010949-10A1 for plasmids, F. Mauri, J. Knöblich, and L. Borsig for cell lines, K. Bruckner, M. Sixt and the Sixt lab for helpful advice, K.B., P. Duchek, K. VijayRaghavan and the Bloomington *Drosophila* Stock Center supported by NIH grant P40OD018537 and the Vienna *Drosophila* Resource Center for fly stocks. E. Ogris for an antibody gift, L. Cooley and S. Munro for contributing the antibodies produced by the Developmental Studies Hybridoma Bank, which was created by the Eunice Kennedy Shriver National Institute of Child Health and Human Development of the NIH, and is maintained at the University of Iowa. We thank the Life Scientific Service Units at IST Austria for technical support and assistance with microscopy and FACS analysis, and J. Friml, C. Guet, P. Rangan, and T. Hurd for comments on the manuscript. K.V. was supported by a DOC fellowship from the Austrian Academy of Sciences. A.G. and A.R. were supported by the Austrian Science Fund (FWF) grant DASI_FWF01_P29638S, D.E.S. by Marie Curie CIG 334077/IRTIM, and A.R. also by Marie Curie *IIF* GA-2012-32950 BB: DICJI. M.R. was supported by the NO Forschungs und Bildungsges.m.b.H. M.M. received funding from the European Union’s Horizon 2020 research and Innovation programme under the Marie Sklodowska-Curie Grant Agreement No. 665385. We are deeply grateful to R. Lehmann in whose lab the work underlying this project began.

## Author contributions

J.B., K.V., M.R., D.E.S., S.E, S.W., H.C., K.S. designed experiments, J.B., M.M., K.V., M.R., I.L., J.S., S.W., S.E., A.G., K.S. carried out experiments, J.B., K.V., M.R., A.R., D.E.S., S.E., H.C., S.W., M.M., K.S. analyzed data, K.V., A.G. and D.E.S. made figures and wrote the manuscript. All authors read and approved the final manuscript.

## MATERIALS AND METHODS

### Fly work

Flies were raised on food bought from IMBA (Vienna, Austria) which contained the standard recipe of agar, cornmeal, and molasses with the addition of 1.5% Nipagin. Adults were placed in cages in a Percival DR36VL incubator maintained at 29°C and 65% humidity; embryos were collected on standard plates prepared in house from apple juice, sugar, agar and Nipagin supplemented with yeast from Lesaffre (Marcq, France) on the plate surface. Embryo collections for fixation (7 hour collection) as well as live imaging (4.5 hour collection) were conducted at 29°C.

### Fly Lines utilized

*srpHemo-GAL4* was provided by K. Brückner (UCSF, USA) (Bruckner et al., 2004), *UAS-CG8602∷FLAG∷HA* (from K. VijayRaghavan National Centre for Biological Sciences, Tata Institute of Fundamental Research) (Guruharsha et al., 2011). The stocks *w*^*1118*^; *minerva*^*3102*^ (BDSC-17262), (*pn*^*1*^;; *ry*^*503*^*Dr*^*1*^P[Δ 2-3] (BDSC-1429), *Df(3L)BSC117* (BDSC-8976), *Oregon R* (BDSC-2375), *w*^*-*^; *P{w[+mC]=UAS-mCherry.NLS}2;MKRS/Tm6b, Tb[1]* (BDSC-38425), *w*^*-*^,*P{UAS-Rab11-GFP}2* (BDSC-8506), *y[1] sc[*] v[1]; P{y[+t7.7] v[+t1.8]=TRiP.GL00069}attP2* (BDSC-35195), *y[1] w[*]; Mi{y[+mDint2]=MIC}GlcAT-P[MI05251]/TM3, Sb[1]* (BDSC-40779) were obtained from the Bloomington *Drosophila* Stock Centre, Bloomington, USA. The RNAi lines v60100, v110406, v2826, v101575 were obtained from the Vienna Drosophila Resource Center (VDRC), Vienna, Austria. Lines *w*^*-*^; *P{w[+mC*; *srpHemo-3xmcherry}, w*^*-*^; *P{w[+mC; srpHemo-H2A∷3xmcherry}* were published previously (Gyoergy et al., 2018).

### Lines used in figures

**Figure 1D-H:** *w-; +; srpHemo-3xmcherry*. **I-K**: *w*^*-*^ *P(w+)UAS-dicer****/****w-; P{attP,y[+],w[3`]/+; srpHemo-Gal4 UAS-GFP, UAS-H2A∷RFP/+, w*^*-*^ *P(w+)UAS-dicer2/w-; RNAi C1GalTA (v110406)/+; srpHemo-Gal4 UAS-GFP, UAS-H2A:RFP/+*. **L**: *w-;* +; *srpHemo-H2A∷3xmcherry, w-; C1GalTA*^*2.1*^; *srpHemo-H2A∷3xmcherry* **M**: *w-; srpHemo-H2A∷3xmcherry, w-; srpHemo-H2A∷3xmcherry, Mi{MIC}GlcAT-PMI05251* **Figure S1A-L**: *w-; +; srpHemo-3xmcherry*. **M, N, P**: *w-, UAS-Dicer2/w-; P{attP,y[+],w[3`]/+; srpHemo-Gal4 UAS-GFP, UAS-H2A∷RFP/+*, *w-UAS-Dicer2/ w-; RNAi C1GalTA (v110406)/+; srpHemo-Gal4 UAS-GFP, UAS-H2A∷RFP/+* **O:** w-; + *srpHemo-H2A∷3xmcherry, w-; C1GalTA*^*2.1*^; *srpHemo-H2A∷3xmcherry* **Q**: *w-; srpHemo-H2A∷3xmcherry, w-; srpHemo-H2A∷3xmcherry, Mi{MIC}GlcAT-PMI05251* **Figure 2A, B, E:** *w-; +; srpHemo-3xmcherry,* **D**: *Oregon R*. **F, G, H:** *w-; +; srpHemo-3xmcherry, w-; +; srpHemo-3xmcherry,P{EP}CG8602*^*3102*^. **I:** *w-; srpHemo-Gal4; UAS-CG8602∷FLAG∷HA* **Figure S2A, B:** *Oregon R*. **D**: *P{EP}CG8602*^*3102*^ **Figure 3A:** *w-; +; srpHemo-H2A∷3xmcherry*. **B:** *w-; +; srpHemo-H2A∷3xmcherry, P{EP}CG8602*^*3102*^. **C:** *w-; srpHemo-CG8602; srpHemo-H2A∷3xmcherry P{EP}CG8602*^*310*2^. **D:** Control: *w-; srpHemo-Gal4 UAS-mcherry∷nls;* +, mutant: *w-; srpHemo-Gal4 UAS-mcherry∷nls; P{EP}CG8602*^*3102*^. Df cross: *w-; srpHemo-Gal4 UAS-mcherry:nls; P{EP}CG8602*^*3102*^*/ Df(3L)BSC117, HA*, rescue: *w-; srpHemo-Gal4 UAS-mcherry:nls; UAS-CG8602∷FLAG∷HA P{EP}CG8602*^*3102*^, precise excision: *srpHemo-Gal4 UAS-mcherry:nls; P{EP}CG8602*^*3102*^*Δ32*. **E:** *w*^*-*^ *P(w+)UAS-dicer****/****+; +; srpHemo-Gal4 UAS-GFP UAS-H2A:RFP/+, w-UAS-dicer2/w-; RNAi CG8602 (v101575)/+; srpHemo-Gal4 UAS-GFP UAS-H2A:RFP/+*. **F**: *w*^*-*^ *P(w+)UAS-dicer****/****y[1] v[1]; srpHemo-Gal4 UAS-mcherry∷nls;* +, *w-; srpHemo-Gal4 UAS-mcherry∷nls; P{EP}CG8602*^*3102*^ **G**: w-; +; srpHemo-3xmcherry, w-; +; srpHemo-3xmcherry *P{EP}CG8602*^*3102*^, w-; shg^P34^; srpHemo-3xmcherry *P{EP}CG8602*^*3102*^ H: w-; +; srpHemo-3xmcherry, w-; +; srpHemo-3xmcherry *P{EP}CG8602*^*3102*^ **I-L**: w-; +; srpHemo-H2A∷3xmcherry, w-; +; srpHemo-H2A∷3xmcherry *P{EP}CG8602*^*3102*^ **Figure S3A:** w-; +; srpHemo-H2A∷3xmcherry, w-; +; srpHemo-H2A∷3xmcherry *P{EP}CG8602*^*3102*^, w-; srp-CG8602; srpHemo-H2A∷3xmcherry *P{EP}CG8602*^*3102*^ **B, C, E:** *w-; +; srpHemo-Gal4 UAS-GFP UAS-H2A:RFP/+, w-; RNAi CG8602 (v101575)/+; srpHemo-Gal4 UAS-GFP UAS-H2A∷RFP/+*. **D, F**: w-; +; srpHemo-H2A∷3xmcherry, w-; +; srpHemo-H2A∷3xmcherry *P{EP}CG8602*^*3102*^ **Figure 4A-H:** *w-; +, srpHemo-3xmcherry, w-; +, srpHemo-3xmcherry P{EP}CG8602*^*3102*^. **Figure 4I, S4D, F** Control: *w/ y,w[1118];*^*-*^; *P{attP,y[+],w[3`]}/srpHemo-Gal4; srpHemo-H2A∷3xmcherry/+;* Qsox1 RNAi: *w*^*-*^ *w/ y,w[1118];*^*-*^; *v108288/srpHemo-Gal4; srpHemo-H2A∷3xmcherry/+*.**. Figure 4J, S4E** w-; *srpHemo-3xmcherry*; w-; *P{SUPor-P}Qsox1KG04615; srpHemo-3xmcherry* **Figure 5C**: *w-; srpHemo-MFSD1; srpHemo-H2A∷3xmcherry P{EP}CG8602*^*3102*^, **F**: *w-; +; srpHemo-3xmcherry, w-; +; srpHemo-3xmcherry P{EP}CG8602*^*3102*^, : *w-; srpHemo-MFSD1; srpHemo-3xmcherry P{EP}CG8602*^*3102*^

### Embryo Fixation and Immunohistochemistry

Embryos were collected on apple juice plates from between 6 and 8.5 hours at 29°C. Embryos were incubated in 50% Chlorox (DanClorix) for 5 min and washed. Embryos were fixed with 17% formaldehyde/heptane for 20 min followed by methanol or ethanol devitellinization except for T antigen analysis, when embryos were fixed in 4% paraformaldehyde/heptane. Fixed embryos were blocked in BBT (0.1M PBS + 0,1% TritonX-100 + 0,1% BSA) for 2 hours at RT. Antibodies were used at the following dilutions: α-T antigen (Steentoft et al., 2011) 1:5, α-GFP (Aves Labs Inc., Tigard, Oregon) 1:500 and incubated overnight at 4°C (GFP) or room temperature (T antigen). Afterwards, embryos were washed in BBT for 2 hours, incubated with secondary antibodies (ThermoFisher Scientific, Waltham, Massachusetts, USA) at RT for 2 hours, and washed again for 2 hours. Vectashield (Vector Laboratories, Burlingame, USA) was then added. After overnight incubation in Vectashield at 4°C, embryos were mounted on a slide and imaged with a Zeiss Inverted LSM700 Confocal Microscope using a Plain-Apochromat 20X/0.8 Air Objective or a Plain-Apochromat 63X/1.4 Oil Objective.

### Lectin staining

Embryos were fixed with 10% formaldehyde/heptane and devitellinized with Ethanol. Blocking was conducted in BBT for 2 hours at room temperature. A FITC-labeled lectin kit #2 (EY laboratories, San Mateo, USA) was utilized (table below summarizes abbrevations of used lectins). Each lectin was diluted to 1:25 and incubated with fixed embryos overnight at room temperature (RT). Embryos were washed in BBT for 2 hours at RT and Vectashield was added. After overnight incubation at 4°C, embryos were mounted on a slide and imaged with a Zeiss Inverted LSM700 Confocal Microscope using a Plain-Apochromat 63X/1.4 Oil Objective. Macrophages in late Stage 11 embryos were imaged at germband entry and evaluated by eye for enriched staining on macrophages compared to other tissues.

|  |  |  |  |  |  |  |
| --- | --- | --- | --- | --- | --- | --- |
| **Lectin** | peanut<br>agglutinin | Ulex<br>europaeus<br>agglutinin | Wheat<br>germ<br>agglutinin | Griffonia<br>simplicifolia<br>agglutinin I | Maclura<br>pomifera<br>agglutinin | Griffonia<br>simplicifolia<br>agglutinin II |
| **Abbreviation** | PNA | UEA-I | WGA | GS-I | MPA | GS-II |
| **Lectin** | Soyabean<br>agglutinin | Dolichos<br>biflorus<br>agglutinin | Concavali<br>n A | Helix<br>pomatia<br>agglutinin | Limulus<br>polyphenus<br>agglutinin | Bauhinia<br>purpurea<br>agglutinin |
| **Abbreviation** | SBA | DBA | ConA | HPA | LPA | BPA |

### In situ hybridization

Embryos were fixed with 10% formaldehyde/heptane for 20 min followed by methanol devitellinization for *in situ* hybridization. A 590bp piece of the CG8602 gene with T7 promoter was synthesized using Fw primer TTCATGTGCCTGCTGGGATT, Rv primer GATAATACGACTCACTATAGGGTTACGCTGCAAAATCCGCT from the whole fly DNA prep (see below). T7 polymerase-synthesized digoxigenin-labelled anti-sense probe preparation and *in situ* hybridization was performed using standard methods (Lehmann and Tautz, 1994). Images were taken with a Nikon-Eclipse Wide field microscope with a 20X 0.5 NA DIC water Immersion Objective.

### Macrophage extraction

Embryos were bleached in 50% Chlorox in water for 5 minutes at RT. Stage late 11/early 12 embryos were lined up and then glued to 50 mm Dish No. 0 Coverslip, 14 mm Glass Diameter, Uncoated dish (Zeiss, Germany). Cells from the germband margin were extracted using a ES Blastocyte Injection Pipet (spiked, 20µm inner diameter, 55mm length; BioMedical Instruments, Germany). Extracted cells were placed in Schneider’s medium (Gibco) supplemented with 20% FBS (Sigma-Aldrich, Saint Louis, Missouri, USA).

### Immunohistochemistry of extracted macrophages

Extracted macrophages were collected by centrifugation at 500g for 5 min at room temperature. The cell pellet was resuspended in a small volume of Phospho-buffered saline (PBS) and smeared on a cover slip. The cell suspension was left to dry before cells were fixed with 4% paraformaldehyde in 0.1M Phosphate Buffer for 20 min at room temperature. Cells were washed 3 times in 0.1M PBS and permeabilized in 0.5% Triton-X 100 in PBS. Cells were blocked for 1 hour at room temperature in 20% Fetal Bovine Serum + 0.25% Triton X-100 in PBS. Primary antibodies were diluted in blocking buffer: anti-HA (Roche, Basel, Switzerland) 1:50, anti-Golgin 84, 1:25, anti-Calnexin 99a 1:25, anti-Hrs.8.2 1:25 or anti-Rab7 1:25 all from DSHB (Riedel et al., 2016), and incubated for 1 hour at room temperature. Cells were then washed 5 times in blocking buffer. Secondary antibodies were diluted in blocking buffer: anti-rat 633 1:300, anti-mouse 488 1:300 (both from ThermoFisher Scientific, Waltham, Massachusetts, USA). Secondary antibodies were incubated for 1 hour at room temperature. Cells were washed 5 times in PBS + 0.1% Triton X-100 and mounted in VectaShield+DAPI (LifeTechnologies, Carlsbad, USA) utilized at 1:75.

### S2 cell work

S2R+ cells (a gift from Frederico Mauri of the Knoblich laboratory at IMBA, Vienna) were grown in Schneider’s medium (Gibco) supplemented with 10% FBS (Gibco) and transfected with PTS1-GFP (a gift from Dr. McNew) and/or the *srpHemo-CG8602∷3xmcherry* construct using Effectene Tranfection Reagent (Qiagen, Hilden, Germany) following the manufacturer’s protocol. Transfected S2R+ cells were grown on Poly-L-Lysine coated coverslips (ThermoFisher Scientific, Waltham, Massachusetts, USA) in complete Schneider’s medium (Gibco) supplemented with 10% FBS (Sigma-Aldrich, Saint Louis, Missouri, USA) and 1% Pen/Strep Gibco() to a confluency of 60%. To visualize lysosomes, cells were incubated with Lysotracker 75nM Green DND-26 (Invitrogen) in complete Schneider’s medium for 30 minutes at 25°C. Cells were washed in complete Schneider’s medium 3 times before imaging on an inverted LSM-700 (Zeiss). To visualize mitochondria, mitotracker Green FM (Invitrogen) was diluted in prewarmed Schneider’s medium supplemented with 1% Pen/Strep to a concentration of 250nM. Cells were incubated in the Mitotracker solution for 45 minutes at 25°C. Cells were then washed 3 times in complete Schneider’s medium before imaging.

### DNA Isolation from Single Flies

Single male flies were frozen for at least 3 hours before grinding them in 100mM Tris-HCl, 100mM EDTA, 100mM NaCL and 0.5% SDS. Lysates were incubated at 65°C for 30 minutes. Then 5M KAc and 6M LiCl were added at a ratio of 1:2.5 and lysates were incubated on ice for 10 min. Lysates were centrifuged for 15 minutes at 20,000xg, supernatant was isolated and mixed with Isopropanol. Lysates were centrifuged again for 15 minutes at 20.000xg, supernatant was discarded and the DNA pellet was washed in 70% EtOH and subsequently dissolved in ddH20.

### FACS sorting

Embryos were collected for 1 hour and aged for an additional 5 hours, all at 29°C. Embryos collected from w-flies were processed in parallel and served as a negative control. Embryos were dissociated as described previously (Gyoergy et al., 2018). The cells were sorted using a FACS Aria III (BD) flow cytometer. Emission filters were 600LP, 610/20 and 502 LP, 510/50. Data was analyzed with FloJo software (Tree Star). The cells from the dissociated negative control *w*^*-*^ embryos were sorted to set a baseline plot.

### qPCR

RNA was isolated from approximately 50,000 mCherry positive or mCherry negative cells using RNeasy Plus Micro Kit (Qiagen, Hilden, Germany following manufacturer’s protocol. Resulting RNA was used for cDNA synthesis using Sensiscript RT Kit (Qiagen, Hilden, Germany) and oligodT primers. A Takyon qPCR Kit (Eurogentec) was used to mix qPCR reactions based on the provided protocol. qPCR was run on a LightCycler 480 (Roche, Basel, Switzerland) and data were analyzed in the LightCycler 480 Software and Prism (GraphPad Software). Data are represented as relative expression to a housekeeping gene (=2-^Δct^) or fold change in expression (=2-^ΔΔct^). Primer sequences utilized for flies were obtained from the FlyPrimerBank (http://www.flyrnai.org/FlyPrimerBank). Minerva/CG8602: Fw pr TGTGCTTCGTGGGAGGTTTC, Rv pr GCAGGCAAAGATCAACTGACC. C1GalTA: Fw pr TGCCAACAGTCTGCTAGGAAG, Rv pr CTGTGATGTGCATCGTTCACG. Ugalt: Fw pr GCAAGGATGCCCAGAAGTTTG, Rv pr GATATAGACCAGCGAGGGGAC. RpL32: Fw pr AGCATACAGGCCCAAGATCG, Rv pr TGTTGTCGATACCCTTGGGC

### Protein preps from embryos for Western

Embryos were collected for 7 hours at 29°C, bleached and hand-picked for the correct Stage. 50-200 embryos were smashed in RIPA buffer (150mM NaCl, 0,5% Sodiumdeoxychalat, 0,1% SDS, 50mM Tris, pH 8) with Protease inhibitor (Complete Mini, EDTA free, Roche, Basel, Switzerland) using a pellet homogenizer (VWR, Radnor, USA) and plastic pestles (VWR, Radnor, USA) and incubated on ice for 30 min.

Afterwards, samples were centrifuged at 4°C, 16,000g for 30 min and the supernatant was collected and used for experiments. The protein concentration was quantified using the Pierce BCA Protein Assay Kit (ThermoFisher Scientific, Waltham, Massachusetts, USA).

### Western Blots

30 µg of protein samples were loaded on a 4-15% Mini-PROTEAN TGX Precast Protein Gel (Bio-Rad, Hercules, USA) and run at 100V for 80 min in 1x running buffer (25mM Tris Base, 190mM glycine and 0.1%SDS) followed by transfer onto Amersham Protran Premium 0.45 µm NC (GE Healthcare Lifescience, Little Chalfont, UK) or Amersham Hybond Low Fluorescence 0.2 µm PVDF (GE Healthcare Lifescience, Little Chalfont, UK) membrane using a wet transfer protocol with 25mM Tris Base, 190 mM Glycine + 20% MeOH at either 100 Volts for 60 min or 200mA for 90 min at Mini Trans-Blot Cell Module (Bio-Rad, Hercules, USA). Membranes were blocked in PBS-T (0.1% Triton X-100 in PBS) containing 2% BSA or Pierce Clear Milk Blocking Buffer (ThermoFisher Scientific, Waltham, Massachusetts, USA) for 1 hour at RT. Primary antibodies were incubated overnight at 4°C at the following concentrations: α-T antigen (Copenhagen) 1:10, α-profilin (Verheyen and Cooley, 1994, DSHB) 1:500, anti-GFP (clone 2B6, Ogris lab, MFPL), anti-myc (clone 4A6, Merck Millipore), anti-mouse MFSD1 (Markus Damme, University Kiel), anti-GAPDH (ab181603, Abcam, Cambridge, UK).

Afterwards, blots were washed 3x for 5 min in blocking solution and incubated with Goat anti Mouse IgG (H/L):HRP (Bio-Rad, Hercules, USA) or goat-anti-rabbit IgG (H+L)-HRP (Bio-Rad, Hercules, USA) at 1:5 000 - 10,000 for 1-2 hours at room temperature. Blots were washed 2x 5 min in blocking solution and 1x 5 min with PBS-T. Blots were developed using SuperSignal West Femto Maximum Sensitivity Substrate (ThermoFisher Scientific, Waltham, Massachusetts, USA) according to manufacturer’s instructions. Chemiluminescent signal was detected using the Amersham Imager 600 (GE Healthcare Lifescience) or VersaDoc (Bio-Rad). Images were processed with ImageJ.

### Time-lapse imaging, tracking, speed and persistence analysis

Embryos were dechorionated in 50% bleach for 5 min, washed with water, and mounted in halocarbon oil 27 (Sigma-Aldrich, Saint Louis, Missouri, USA) between a coverslip and an oxygen permeable membrane (YSI). The anterior dorsolateral region of the embryo was imaged on an inverted multiphoton microscope (TrimScope II, LaVision) equipped with a W Plan-Apochromat 40X/1.4 oil immersion objective (Olympus). mCherry was imaged at 1100 nm excitation wavelengths, using a Ti-Sapphire femtosecond laser system (Coherent Chameleon Ultra) combined with optical parametric oscillator technology (Coherent Chameleon Compact OPO). Excitation intensity profiles were adjusted to tissue penetration depth and Z-sectioning for imaging was set at 1µm for tracking and segmentation respectively. For long-term imaging, movies were acquired for 132 - 277 min with a frame rate of 40 sec. All embryos were imaged with a temperature control unit set to 28.5°C.

Images acquired from multiphoton microscopy were initially processed with InSpector software (LaVision Bio Tec) to compile channels from the imaging data, and the exported files were further processed using Imaris software (Bitplane) to visualize the recorded channels in 3D. Macrophage speed and persistence were calculated by using embryos in which the macrophage nuclei were labeled with *srpHemo-H2A∷3XmCherry* (Gyoergy et al., 2018). The movie from each imaged embryo was rotated and aligned along the AP axis for tracking analysis. Increasing the gain allowed determination of germband position from the autofluorescence of the yolk. Movies for vnc analysis were analyzed for 2 hours from the time point that cells started to dive into the channels to reach the outer vnc. Macrophage nuclei were extracted using the spot detection function and nuclei positions in xyz-dimensions were determined for each time point and used for further quantitative analysis. Cell speeds and directionalities were calculated in Matlab (The MathWorks Inc., Natick, Massachusetts, USA) from single cell positions in 3D for each time frame measured in Imaris (Bitplane). Instantaneous velocities from single cell trajectories were averaged to obtain a mean instantaneous velocity value over the course of measurement. To calculate directionality values, single cell trajectories were split into segments of equal length (10 frames) and calculated via a sliding window as the ratio of the distance between the macrophage start-to-end location over the entire summed distance covered by the macrophage between successive frames in a segment. Calculated directionality values were averaged over all segments in a single trajectory and all trajectories were averaged to obtain a mean directionality value for the duration of measurement, with 0 being the lowest and 1 the maximum directionality.

### Fixed embryo image analysis for T antigen levels

Embryos were imaged with a 63x Objective on a Zeiss LSM700 inverted. 10µm stacks (0.5µm intervals) were taken for properly staged and oriented embryos, starting 10µm deep in the tissue. These images were converted into Z-stacks in Fiji. ROIs were drawn around macrophages (signal), copied to tissue close by without macrophages (background) and the average intensity in the green channel of each ROI was measured. For each pair of ROIs background was subtracted from signal individually. The average signal from control ROIs from one imaging day and staining was calculated and all data point from control, mutant and rescue from the same set was divided by this value. This way we introduced an artificial value called Arbitrary Unit (AU) that makes it possible to compare all the data with each other, even if they come from different imaging days when the imaging laser may have a different strength or from different sets of staining. Analysis was done on anonymized samples.

### Macrophage cell counting

Transmitted light images of the embryos were used to measure the position of the germband to determine the stages for analysis. The extent of germband retraction away from the anterior along with the presence of segmentation was used to classify embryos. Embryos with germband retraction of between 29-31% were assigned to late Stage 11. Those with 29-41% retraction for all experiments except the *punt* RNAi (Fig 4J) in which 35-45% was used (both early Stage 12) were analyzed for the number of macrophages that had entered the germband and those with 50-75% retraction (late Stage 12) for the number along the ventral nerve cord (vnc), and in the whole embryo. Macrophages were visualized using confocal microscopy with a Z-resolution of 3µm and the number of macrophages within the germband or the segments of vnc was calculated in individual slices (and then aggregated) using the Cell Counter plugin in FIJI.

To check that this staging allows embryos from the control and *mrva*^*3102*^ mutant to be from the same time during development, embryos were collected for 30 minutes and then imaged for a further 10 hours using a Nikon-Eclipse Wide field microscope with a 20X 0.5 NA DIC water Immersion Objective. Bright field images were taken every 5 minutes, and the timing of the start of the movies was aligned based on when cellularization occurred. We found no significant difference in when germband retraction begins (269.6±9 min in control and 267.1±3 min in *mrva*^*3102*^, p=0.75) or in when the germband retracts to 41% (300±9 min for control, 311±5 min in *mrva*^*3102*^, p=0.23), or in when the germband retraction is complete (386.5±10 min for control, 401.6±8 min for *mrva*^*3102*^, p=0.75). n=10 embryos for control and 25 embryos for *mrva*^*3102*^.

### Cloning

Standard molecular biology methods were used and all constructs were sequenced by Eurofins before injection into flies. Restriction enzymes *BSi*WI, and *Asc*I were obtained from New England Biolabs, Ipswich, Massasuchetts, USA (Frankfurt, Germany). PCR amplifications were performed with GoTaq G2 DNA polymerase (Promega, Madison, USA) using a peqSTAR 2X PCR machine from PEQLAB, (Erlangen, Germany). All Infusion cloning was conducted using an Infusion HD Cloning kit obtained from Clontech’s European distributor (see above); relevant oligos were chosen using the Infusion primer Tool at the Clontech website.

### Construction of *srpHemo-*minerva

A 1467 bp fragment containing the Minerva (CG8602) ORF was amplified from the UAS-CG8602:FLAG:HA construct (DGRC) using primers Fw GAAGCTTCTGCAAGGATGGCGCGCGAGGACGAGGAAC, Rv CGGTGCCTAGGCGCGCTATTCAAAGTTCTGATAATTCTCG. The fragment was cloned into the srpHemo plasmid (a gift from Katja Brückner, (Bruckner et al., 2004)) after its linearization with AscI, using an Infusion HD cloning kit.

### Construction of *srpHemo-MFSD1*

A 1765 bp fragment containing the MFSD1 ORF was amplified from cDNA prepared from dendritic cells (a gift from M. Sixt lab) with Fw primer TAGAAGCTTCTGCAACTTTGCTTCCTGCTCCGTTC, Rv primer ATGTGCCTAGGCGCGAAGGAAAGGCTTCATCCGCA). The fragment was cloned into the srpHemo plasmid (a gift from Katja Brückner, (Bruckner et al., 2004)]) using an Infusion HD cloning kit (Clontech) after its linearization with AscI (NEB).

### Construction of *srpHemo-mrva∷3xmCherry*

Minerva (CG8602) was amplified from a DNA prep from Oregon flies (Fw primer: AGAGAAGCTTCGTACGCGACAACCCTGCTCTACAGAG; Rv primer CGACCTGCAGCGTACGACCCGATCCTTCAAAGTTCTG). The vector, PCasper4 containing a 3xmCherry construct under the control of the srpHemo promoter (Gyoergy et al., 2018), was digested with BsiWI according to the manufacturer’s protocol. The vector and insert were homologously recombined using the In-Fusion HD Cloning Kit. **Generation of pInducer20-MFSD1-eGFP constructs:** For C-terminal tagging MFSD1 was PCR amplified from cDNA prepared from dendritic cells (a gift from M. Sixt lab) with the following primers; fw: GATCTCGAGATGGAGGACGAGGATG; rev: CGACCGGTAACTCTGGATGAGAGAGC and digested with XhoI and AgeI (both New England Biolabs, Ipswich, Massasuchetts, USA). This MFSD1 fragment was cloned into XhoI/AgeI digested peGFP-N1 (Addgene, Cambridge, Massachusetts, USA). C-terminally eGFP tagged MFSD1 was further PCR amplified with following primers; fw: GGGGACAAGTTTGTACAAAAAAGCAGGCTTAATGGAGGACGAGGAT; rev: GGGGACCACTTTGTACAAGAAAGCTGGGTATTACTTGTACAGCTC. This fragment was cloned using Gateway BP Clonase II Enzyme mix and Gateway LR Clonase II Enzyme Mix (ThermoFisher Scientific, Waltham, Massachusetts, USA) via donor vector pDonR211 into the final Doxycyclin inducible expression vector pInducer20 (Meerbrey et al., 2011)according to manufacturer’s instructions. pInducer20-MFSD1-eGFP was amplified in stbl3 bacteria (ThermoFisher Scientific, Waltham, Massachusetts, USA).

### Precise excision

*mrva*^*3102*^ flies which contain the 3102 P element insert in the 5’ region of CG8602 were crossed to a line expressing transposase (BL-1429 (*pn*^*1*^; *ry*^*503*^*Dr*^*1*^P[Δ 2-3]). To allow excision of the P Element, males from the F1 generation containing both the P element and the transposase, were crossed to virgins with the genotype Sp/Cyo; PrDr/TM3Ser (gift from Lehmann lab). In the F2 generation white eyed males were picked and singly crossed to Sp/Cyo; PrDr/TM3Ser virgins.

### Mammalian cell culture

MC-38 colon carcinoma cells (gift from Borsig lab) were kept in DMEM supplemented with 10% FCS (Sigma-Aldrich, Saint Louis, Missouri, USA) and Na-Pyruvate (ThermoFisher Scientific, Waltham, Massachusetts, USA). All cells were kept in a humidified incubator at 37°C with 5% CO2. MC-38 cells were transfected with pInducer20-MFSD1-tagged constructs according to the manufacturer’s instructions using Lipofectamin 2000 (ThermoFisher Scientific, Waltham, Massachusetts, USA). Expression of tagged MFSD1 was induced with 100ng/µl of Doxycycline for 24 hours prior subsequent analysis.

### Cell lysis

Cells were lysed in lysis buffer (25mM Tris, 150mM NaCl, 1mM EDTA, 1% Triton X-100) supplemented with protease inhibitor cocktail (Complete, Roche, Basel, Switzerland) for 20 min on ice, followed by centrifugation at 14,000x g, 4°C for 5 min. The protein lysates were stored at −80°C. Protein concentration was determined with the Pierce BCA Protein Assay Kit (ThermoFisher Scientific, Waltham, Massachusetts, USA).

### Immunofluorescence

Cells were fixed with 4% formaldehyde (ThermoFisher Scientific, Waltham, Massachusetts, USA) in PBS for 15 minutes at room-temperature. Cells were washed three times with PBS followed by blocking and permeabilization with 1% BSA (Sigma-Aldrich, Saint Louis, Missouri, USA)/0.3% Triton X-100 in PBS for 1 hour. Antibodies were diluted in blocking/permeabilization buffer and incubated for 2 hours at room temperature. Primary antibodies used were: anti-GFP (clone 5G4, Ogris lab, MFPL), anti-giantin (Biolegend, #19243), anti-GRASP65 (ThermoFisher Scientific, Waltham, Massachusetts, USA, PA3-910), anti-LAMP1 (#ab24170, Abcam, Cambridge, UK), anti-Rab7 (Cell Signalling Technology, #D95F2), anti-Rab5 (Cell Signalling Technology, #C8B1). Cells were washed three times with PBS-Tween20 (0.05%) for 5 minutes each, followed by secondary antibody incubation in blocking/permeabilization buffer for 1 hour at room-temperature. Secondary antibodies used were: goat anti-mouse IgG (H+L) Alexa Fluor 488 (ThermoFisher Scientific, Waltham, Massachusetts, USA, A11001), goat anti-rabbit IgG (H+L) Alexa Fluor 555 (ThermoFisher Scientific, Waltham, Massachusetts, USA, A21428), goat anti-rabbit IgG (H+L) Alexa Fluor 633 (ThermoFisher Scientific, Waltham, Massachusetts, USA, A21070). Cells were counterstained with DAPI (ThermoFisher Scientific, Waltham, Massachusetts, USA) for 10 minutes in PBS-Tween20%. Cells were mounted with ProLong Gold Antifade Mountant (ThermoFisher Scientific, Waltham, Massachusetts, USA, #P36930). Images were acquired using a Zeiss LSM880 confocal microscope. Pictures were processed with ImageJ.

### Embryonic Protein Prep for Glycoproteomics

150 mg fly embryos were homogenized in 2 ml 0.1% RapiGest, 50mM ammonium bicarbonate using a dounce homogenizer. The lysed material was left on ice for 40 min with occasional vortexing followed by probe sonication (5 sec sonication, 5 sec pause, 6 cycles at 60% amplitude). The lysate was cleared by centrifugation (1,000× g for 10 min). The cleared lysate was heated at 80°C, 10 min followed by reduction with 5mM dithiothreitol (DTT) at 60°C, 30 min and alkylation with 10mM iodoacetamide at room temperature (RT) for 30 min before overnight (ON) digestion at 37°C with 25µg trypsin (Roche). The tryptic digests were labeled with dimethyl stable isotopes as described (Boersema et al., 2009). The digests were acidified with 12µL trifluoroacetic acid (TFA), 37°C, 20 min and cleared by centrifugation at 10,000g, 10 min. The cleared acidified digests were loaded onto equilibrated SepPak C18 cartridges (Waters) followed by 3× CV 0.1% TFA wash. Digests were labeled on column by adding 5 mL 30 mM NaBH_3_CN and 0.2% formaldehyde (COH_2_) in 50mM sodium phosphate buffer pH 7.5 (Light, *mrva*^*3102*^), or 30mM NaBH_3_CN and 0.2% deuterated formaldehyde (COD_2_) in 50mM sodium phosphate buffer pH 7.5 (Medium, control). Columns were washed using 3 CV 0.1% FA and eluted with 0.5 mL 50% MeOH in 0.1% FA. The eluates were mixed in 1:1 ratio, concentrated by evaporation, and resuspended in Jacalin loading buffer (175mM Tris-HCl, pH 7.4) Glycopeptides were separated from non-glycosylated peptides by Lectin Weak Affinity Chromatography (LWAC) using a 2.8 m column packed in-house with Jacalin-conjugated agarose beads. The column was washed with 10 CVs Jacalin loading buffer (100 µL/min) before elution with Jacalin elution buffer (175mM Tris-HCl, pH 7.4, 0.8M galactose) 4 CVs, 1 mL fractions. The glycopeptide-containing fractions were purified by in-house packed Stage tips (Empore disk-C18, 3M).

### Quantitative O-glycoproteomic Strategy

The glycopeptide quantification based on M/L isotope labeled doublet ratios was evaluated to estimate a meaningful cut-off ratio for substantial changes (Schjoldager et al., 2015). The labeled glycopeptides produced doublets with varying ratios of the isotopic ions as well as a significant number of single precursor ions without evidence of ion pairs. Labeled samples from control *srpHemo-3xmcherry* embryos and *mrva*^*3102*^ *srpHemo-3xmcherry* mutant embryos were mixed 1:1 and subjected to LWAC glycopeptide enrichment. The distribution of labeled peptides from the LWAC flow-through showed that the quantitated peptide M/L ratios were normally distributed with 99.7% falling within +/-0.55 (Log_10_). We selected doublets with less/more than +/-0.55(Log_10_) value as candidates for isoform-specific O-glycosylation events.

### Mass spectrometry

EASY-nLC 1000 UHPLC (Thermo Scientific) interfaced via nanoSpray Flex ion source to an-Orbitrap Fusion mass spectrometer (Thermo Scientific) was used for the glycoproteomic study. A precursor MS1 scan (m/z 350–1,700) of intact peptides was acquired in the Orbitrap at a nominal resolution setting of 120,000. The five most abundant multiply charged precursor ions in the MS1 spectrum at a minimum MS1 signal threshold of 50,000 were triggered for sequential Orbitrap HCD-MS2 and ETD-MS2 (m/z of 100–2,000). MS2 spectra were acquired at a resolution of 50,000. Activation times were 30 and 200 ms for HCD and ETD fragmentation, respectively; isolation width was 4 mass units, and 1 microscan was collected for each spectrum. Automatic gain control targets were 1,000,000 ions for Orbitrap MS1 and 100,000 for MS2 scans. Supplemental activation (20 %) of the charge-reduced species was used in the ETD analysis to improve fragmentation. Dynamic exclusion for 60 s was used to prevent repeated analysis of the same components. Polysiloxane ions at *m/z* 445.12003 were used as a lock mass in all runs. The mass spectrometry glycoproteomics data have been deposited to the ProteomeXchange Consortium (Vizcaino et al., 2016) via the PRIDE partner repository with the dataset identifier PXD011045.

### Mass spectrometry Data analysis

Data processing was performed using Proteome Discoverer 1.4 software (Thermo Scientific) using Sequest HT Node as previously described (Schjoldager et al., 2015). Briefly, all spectra were initially searched with full cleavage specificity, filtered according to the confidence level (medium, low and unassigned) and further searched with the semi-specific enzymatic cleavage. In all cases the precursor mass tolerance was set to 6 ppm and fragment ion mass tolerance to 20 mmu. Carbamidomethylation on cysteine residues was used as a fixed modification. Methionine oxidation and HexNAc attachment to serine, threonine and tyrosine were used as variable modifications for ETD MS2. All HCD MS2 were pre-processed as described (2) and searched under the same conditions mentioned above using only methionine oxidation as variable modification. All spectra were searched against a concatenated forward/reverse *Drosophila melanogaster*-specific database (UniProt, March 2018, containing 39034 entries with 3494 canonical reviewed entries) using a target false discovery rate (FDR) of 1%. FDR was calculated using target decoy PSM validator node. The resulting list was filtered to include only peptides with glycosylation as a modification.

Glycopeptide M/L ratios were determined using dimethyl 2plex method as previously described (Schjoldager et al., 2015)

### Statistics and Repeatability

Statistical tests as well as the number of embryos/ cells assessed are listed in the figure legends. All statistical analyses were performed using GraphPad Prism and significance was determined using a 95% confidence interval. Data points from individual experiments / embryos were pooled to estimate mean and standard error of the mean. Sample size refers to biological replicates. No statistical method was used to predetermine sample size and the experiments were not randomized. For major questions, data were collected and analyzed masked. Normality was evaluated by D’Agostino & Pearson or Shapiro-Wilk normality test. Unpaired t-test or Mann-Whitney test was used to calculate the significance in differences between two groups and One-Way Anova followed by Tukey post-test or Kruskal-Wallis test followed by Conover or Dunn’s post-test for multiple comparisons.

All measurements were performed in 3-38 embryos. Representative images shown in Fig 1E, F, G, I, Fig 2G, I Fig3 A, B, C Fig 5B, C, F and Supplementary Figures FigS2E-L and FigS5 C,D were from separate experiments repeated 3 to 6 times. FigS1A-M is from separate experiments that were repeated at least twice. Representative *in situ* images shown in Fig 2D and Fig S2A, B, D were from an experiment repeated 3 times. Stills shown in Fig 3I, K and Fig S3H are representative images from two-photon movies, which were repeated at least 3 times.

## Supplementary Material Legends

**Video 1, related to Fig 3: Representative movie of macrophage migration into the germband in the control.** Macrophages (red) are labeled with *srpHemo-H2A∷3xmcherry*. The time interval between each acquisition is 40 sec and the display rate is 15 frames/sec. Scale bar represents 30µm.

**Video 2, related to Fig 3**: **Representative movie of macrophage migration into the germband in the *mrva*^*3102*^ mutant**. Macrophages (red) are labeled with *srpHemo-H2A∷3xmcherry*. The time interval between each acquisition is 40 sec and the display rate is 15 frames/ sec. Scale bar represents 30µm.

**Video 3, related to Fig 3: Representative movie of macrophage migration on the vnc in the control**. Macrophages (red) are labeled with *srpHemo-H2A∷3xmcherry*. The time interval between each acquisition is 40 ec and the display rate is 15 frames/sec. Scale bar represents 30µm

**Video 4, related to Fig 3: Representative movie of macrophage migration on the vnc in the *mrva*^*3102*^ mutant**. Macrophages (red) are labeled with *srpHemo-H2A∷3xmcherry*. The time interval between each acquisition is 40 sec and the display rate is 15 frames/ sec. Scale bar represents 30µm.

**Table S1, related to Fig 4: Mass spectrometric analysis of the T and Tn antigen containing O-glycoproteome from wild type and *mrva*^*3102*^ mutant Stage 11-12 *Drosophila melanogaster* embryos.** Each row lists an individually identified tryptically processed peptide. The 2^nd^–4^th^ columns describe the analyzed peptide. The 5^th^, 6^th^, 7^th^ and 12^th^ are the names and accessions to Uniprot. The 8^th^ indicates the position of the modified amino acid. The 9^th^ indicates the number and 10^th^ the type of glycosylation. The 11^th^ lists the exact position and the 13^th^ the exact description of glycosylation. The 14^th^ is the ratio of the amount of the particular glycopeptide in the control samples (medium) over the amount in the *mrva*^*3102*^ (light). The 15^th^ is the number of missed cleavages after the tryptic digest. The 16^th^ is the measured intensity. The 17^th^ column shows the mass to charge ratio.

**Table S2, related to Fig 4: All candidate proteins with at least 3-fold changes in T and Tn antigen.** Columns list the gene name, the predicted or known function of the gene, if other T or Tn glycosites on the protein are unchanged or changed in the opposite direction, any known human ortholog (identified by BLAST), references for links to cancer and cancer invasion for the mammalian orthologs, the precise site altered, the T and Tn antigen changes observed at a particular glycosylation site, the number of glycosites on the peptide, the peptide sequence and if the glycosylation site is conserved. The site is considered conserved if the human ortholog has a serine or threonine +/-5 amino acids from the *Drosophila* glycosite. References: 1. (Gohrig et al., 2014); 2. (Fan et al., 2018); 3. (Webb et al., 1999); 4. (C.-C. Chiu et al., 2011); 5. (Huang et al., 2016); 6. (Matos et al., 2015); 7. (Cawthorn et al., 2012); 8. (Cao et al., 2015) 9. (Walls et al., 2017); 10.(Zhou et al., 2017); 11. (Linton et al., 2008); 12. (Bian et al., 2016;) 13. (Zhang et al., 2016); 14. (Gonias et al., 2017); 15. (Katchman et al., 2013, 2011); 16. (Stojadinovic et al., 2007); 17. (Zhou et al., 2016); 18. (Hu et al., 2018); 19. (Li et al., 2008); 20. (Senanayake et al., 2012); 21. (Sheu et al., 2014) (Sheu et al., 2014); 22. (Mao et al., 2018); 23.(Yokdang et al., 2016).

**Figure S1.**
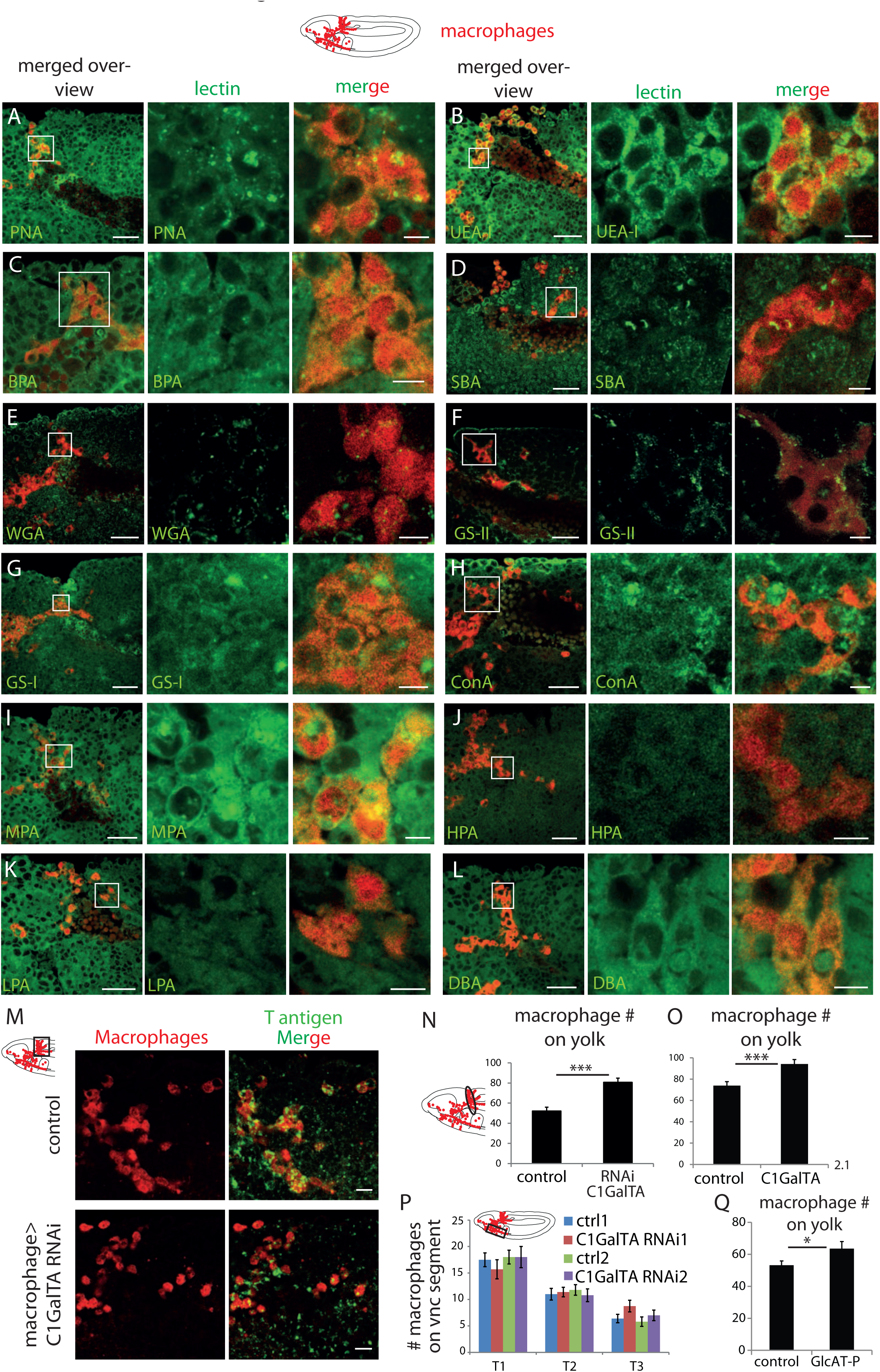
Related to Figure 1: Lectin screen reveals enriched staining for PNA and UEA-1 on macrophages. **(A-L)** Confocal images of fixed late Stage 11/ early Stage 12 wild type embryos schematic above) stained with different lectins (visualized in green) indicated in green type in the lower left corner. Macrophages are detected through srpHemo-3xmCherry expression (red). Boxed area in schematic shows area of merged overview image at left. Boxed area in merged overview corresponds to the images shown magnified at right. **(M)** Confocal images of the germband from fixed early Stage 12 embryos from the control and ones in which UAS-C1GalTA RNAi is expressed in macrophages under srpHemo-GAL4 control. Macrophages visualized with an antibody against GFP expressed in macrophages (srpHemo>GFP) (red) and T antigen by antibody staining (green). Boxed area in schematic at left indicates embryo region imaged. **(N,O)** Quantification of macrophages on the yolk in fixed early Stage 12 embryos in **(N)** srpHemo>UAS-C1GalTA RNAi (vdrc 2826) and **(O)** the C1GalTA[2.1] excision mutant shows an increase in both compared to the control (n=14-24, p=0.00004 for N, p=0.0007 for O). **(P)** Quantification of macrophage number in the vnc segments shown in the schematic in fixed mid Stage 12 embryos detects no difference between control and srpHemo>UAS-C1GalTA RNAi embryos (n=10-20). **(Q)** Quantification of macrophages on the yolk in fixed early Stage 12 embryos in GlcAT-PMI05251 shows a 20% increase compared to the control (n=17-20, p=0.04). Significance was assessed by Mann-Whitney test in **N** and Student’s t-test in **O-Q,** ns=p>0.05, *=p<0.05, ***=p<0.001. Scale bars are 30µm in overview images and 5µm in magnifications in **A-L**, 10µm in **M**.

**Figure S2.**
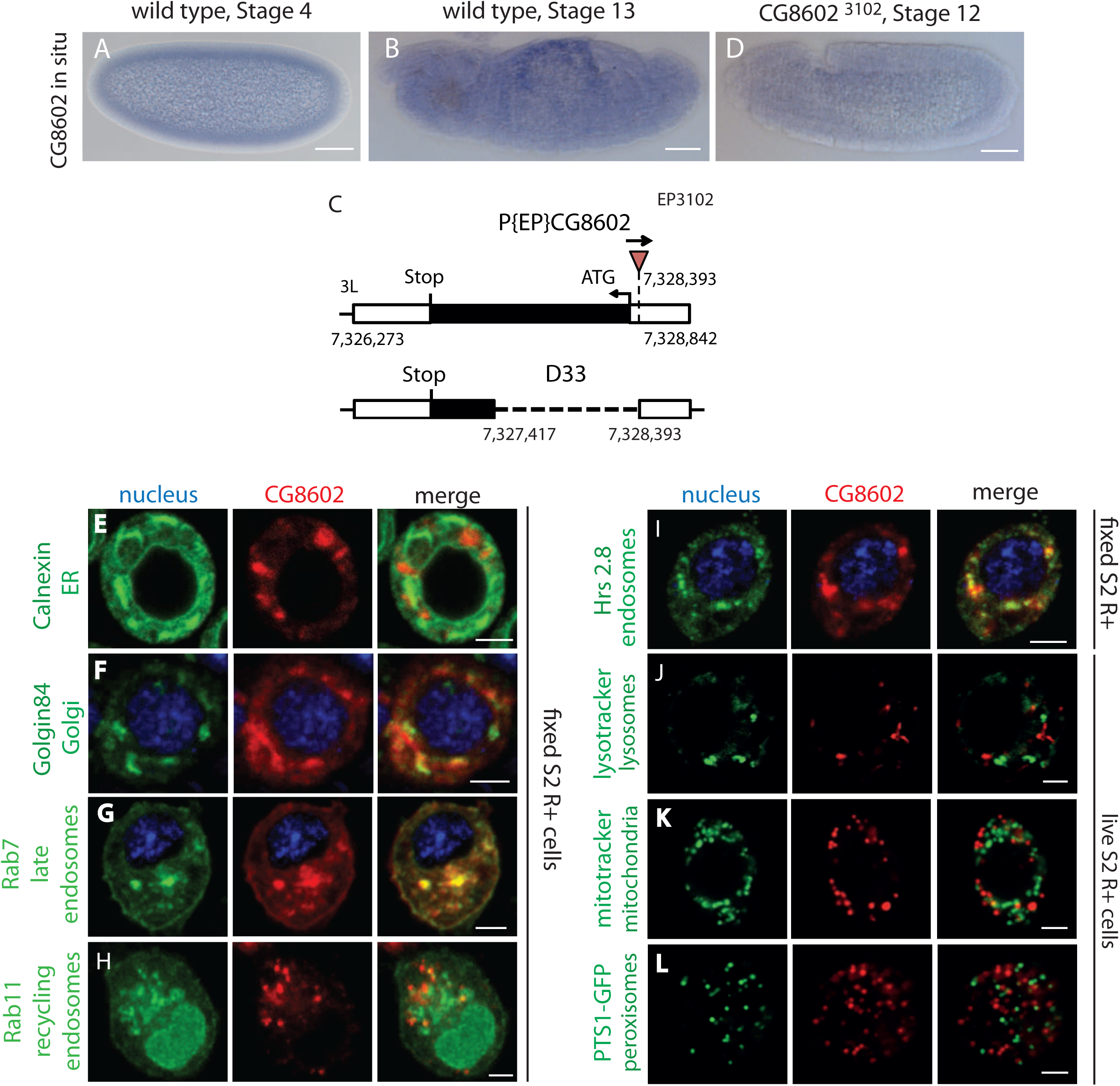
Related to Figure 2: CG8602 expression and localization. **(A-B, D)** *In situ* hybridization of RNA probes against CG8602. In wild type embryos (**A**) maternally deposited CG8602 RNA is evident in Stage 4 embryos and (**B**) uniform lower level expression in Stage 13 embryo, with enrichment in the amnioserosa, but none in macrophages. **(C)** Schematic depicting the CG8602 gene and the insertion site of the EP3102 P element and the Δ33 excision mutant induced by P element mobilization which removes 914 bp of the ORF. (**D)** Expression of CG8602 RNA is strongly reduced in Stage 12 *CG86023102* mutant embryos. (**E-L**) Confocal images of S2R+ cells transfected with (**E-G**) *MT-CG8602∷FLAG∷HA* visualized by HA antibody staining (red) or (**H-L**) *srpHemo-CG8602∷3xmCherry* with different parts of the endomembrane system visualized by antibody staining as indicated (green). DAPI (blue) marks the nucleus. CG8602 showed (**E**) no colocalization with the ER marker Calnexin, partial colocalization with the (**F**) Golgi marker Golgin84, (**G**) late endosomal marker Rab7, (**H**) recycling endosome marker Rab11-YFP, and (**I**) endosomal marker Hrs8.2, no colocalization with (**J**) lysosome marker lysotracker, (**K**) mitochondrial marker mitotracker and (**L**) peroxisomal marker PTS1-GFP in fixed (**E-I**) or live (**J-L**) S2R+ cells. Scale bar is 50μm in A, B and D, 3μm in E-L.

**Figure S3.**
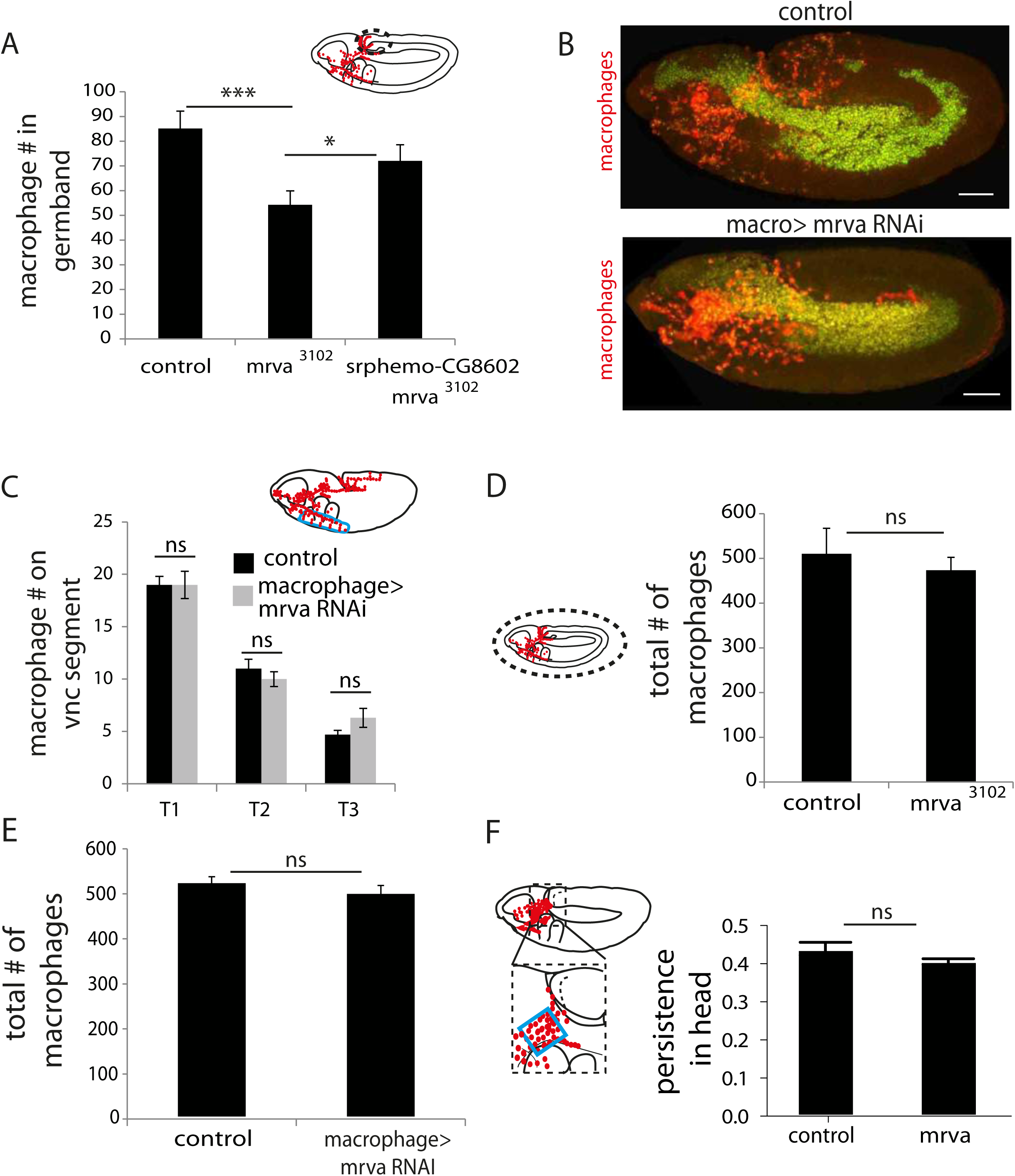
2Related to Figure 3: CG8602 (Minerva) and C1GalTA affect migration into the germband but not along the vnc. **(A)** Quantification of the number of macrophages in the germband in embryos from control, *CG86023102*, and *CG86023102 srpHemo(macro)-CG8602∷HA* showing CG8602 is required in macrophages for invasion of the germband. Macrophages visualized by *srpHemo-H2A∷3xmCherry*. (**B**) Representative confocal images of early Stage 12 embryos from control and *srpHemo(macro)-Gal4* driving *UAS-minerva RNAi* (v101575) expression in macrophages labeled by H2A-RFP (green) and cytoplasmic GFP (red). (**C**) Quantification of the number of macrophages in vnc segments reveals no significant difference in macrophage migration along the vnc between control embryos and those expressing an RNAi against CG8602 (v101575) in macrophages under *srpHemo(macro)-GAL4* control (n=19-20, p>0.05). (**D, E**) Quantification of the total number of macrophages visualized with (**D**) *srpHemo>mcherry∷nls* or (**E**) *srpHemo>H2A∷RFP, GFP* reveals no significant difference between (**D**) control and *CG86023102* mutant embryos (n=15, p>0.05) and (**E**) control and *srpHemo(macro)>CG8602 RNAi* embryos (n=26, p=0.1439). The area analyzed is indicated with the black box in the schematic above**. (F-I)** Quantification of persistence in the head from 2-photon movies with *srpHemo-H2A∷3xmCherry* labeling macrophages shows no change in the *mrva3102* compared to the control. n=3. # tracks: control=329, mutant=340, p=0.2182. (**G**) Quantification of macrophage directionality in the inner vnc shows no change in the *mrva3102* compared to the control n=2,3. # tracks: control=181, mutant=181, p=0.8826. (**I**) Stills at 0, 60 and 120 min reveal no change in macrophage migration in inner vnc in the *mrva3102* mutant compared to the control. Significance was assessed by One-way Anova in A and Student’s t-test in C-F. ns=p>0.05, * p<0.05, *** p<0.001. Scale bars are 50μm in B, 30μm in I.

**Figure S4.**
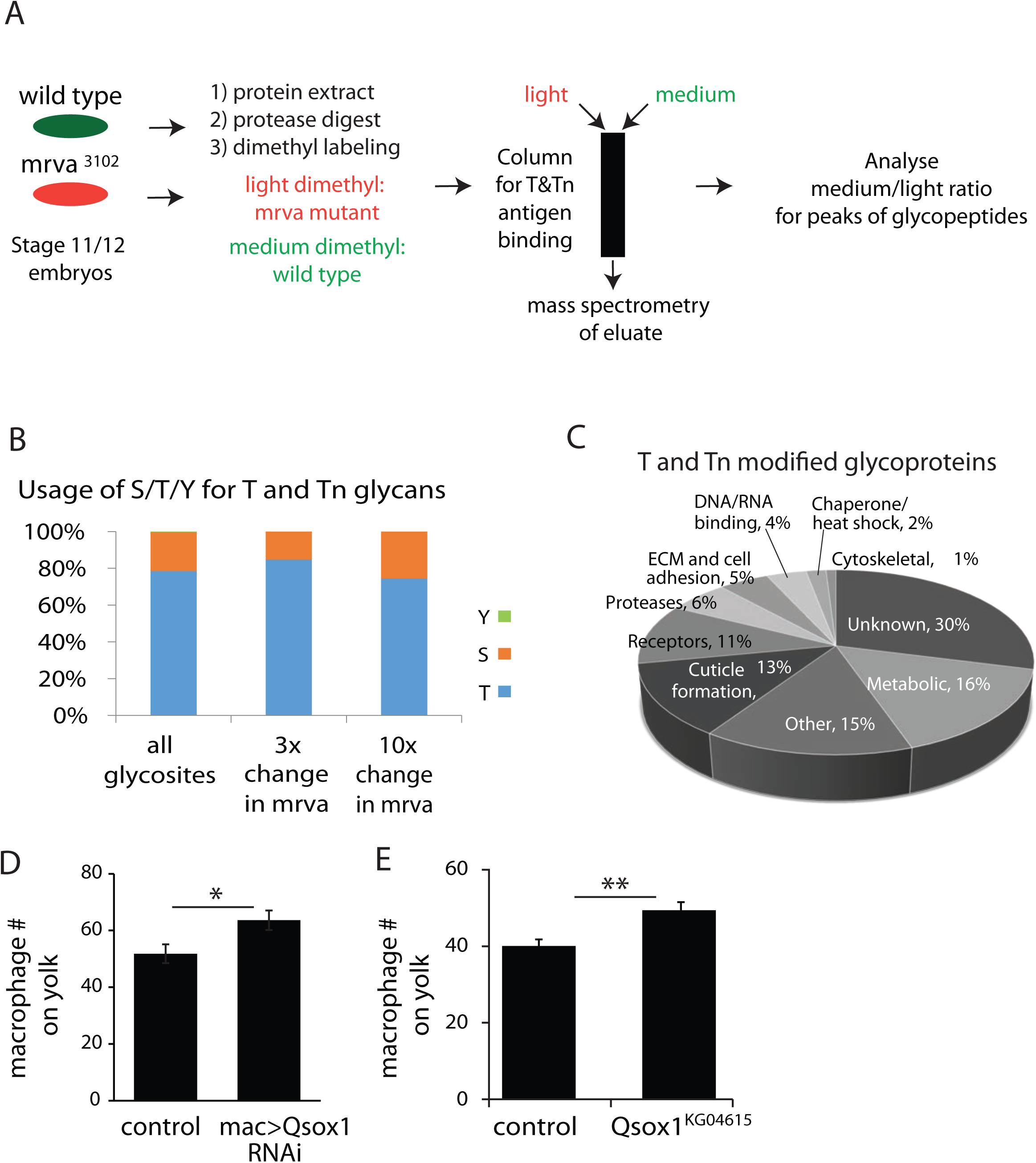
Related to Figure 4 and table 1. **(A)** Work flow for mass spectrometry analysis of T and Tn antigen modification on proteins in stage 11/12 control and mrva^3102^ mutant embryos. **(B)** Similar usage of serine (S), threonine (T) and tyrosine (Y) for glycosylation in all modified proteins in the control and at glycosites that showed at least 3fold and 10fold changes in the mrva^3102^ mutant.**(C)** Analysis of the fractional representation of various functions among all T and Tn antigen modified glycoproteins. **(D)** Increased numbers of macrophages are observed on the yolk neighboring the germband upon knockdown with RNAi v108288 of Qsox1 driven in macrophages by srpHemo-Gal4 (p=0.02) and **(E)** in the full Qsox1 P element (KG04615) mutant compared to the srpHemo-3xmcherry control (p=0.0018). n=24 and 23 for control and RNAi, n=18 for both control and P element mutant. Analyzed by Student’s t test.

**Figure S5.**
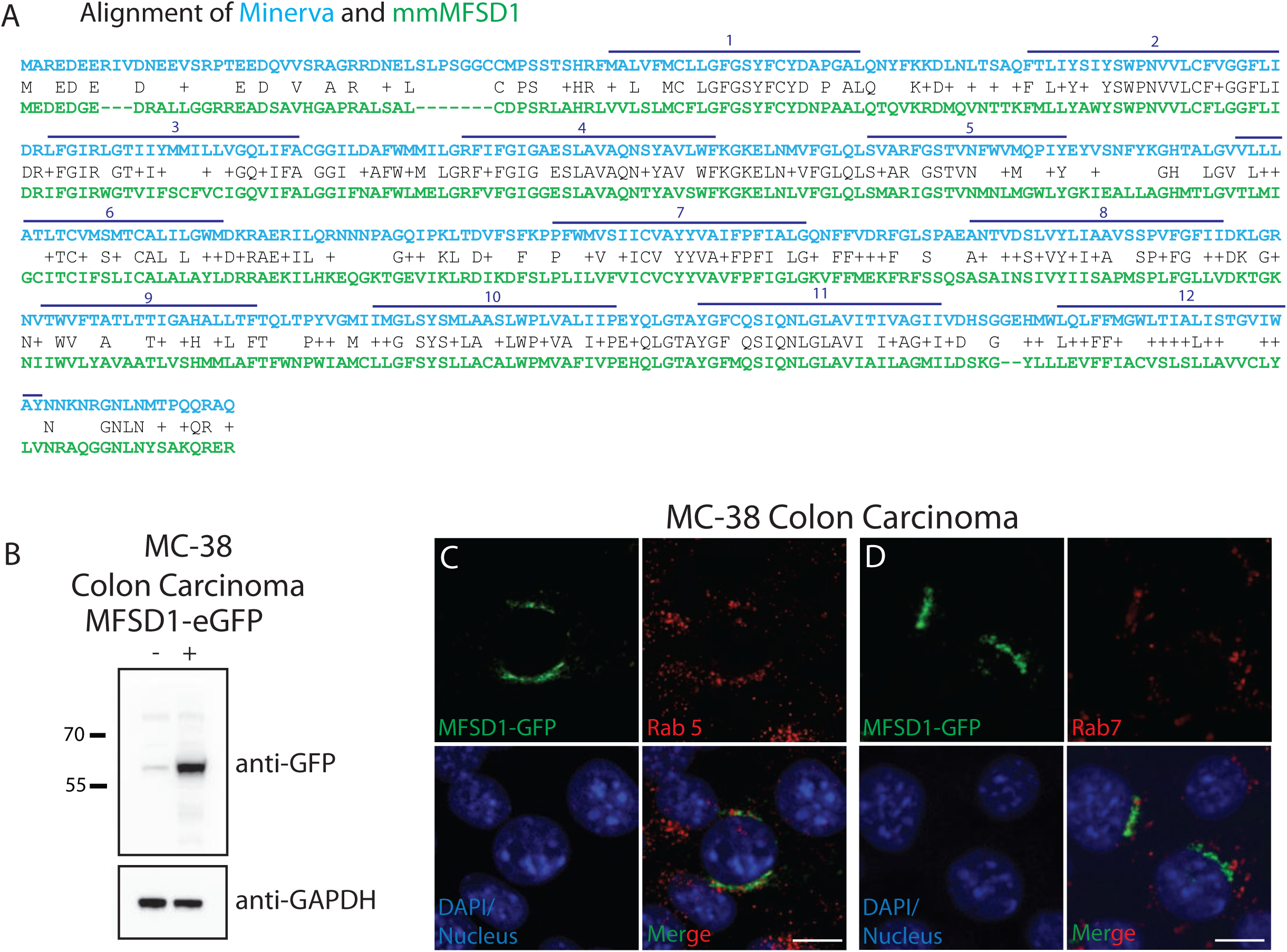
Related to Figure 5: MFSD1-eGFP localization in colon carcinoma. **(A)** Alignment of Minerva and mmMFSD1 by BLAST. The first row in blue type shows the minerva sequence, the second in black identical (one letter symbol) or similar (+) amino acids, and the third in green the mmMFSD1 sequence. Gaps are marked with ‘-‘. The predicted twelve transmembrane domains of Minerva are shown with dark blue lines and numbered above**. (B)** Western blot of MC-38 colon carcinoma cells with (+) and without (-) the induction of MFSD1-eGFP expression from a lentiviraltransduced vector. MFSD1-eGFP was detected with an anti-GFP antibody. GAPDH serves as a loading control. **(C,D)** Co-immunofluorescence of mouse MFSD1-eGFP (green) and **(C)** early endosome marker Rab5 (red) or **(D)** late endosomes marker Rab7 (red) in MC-38 colon carcinoma cells show little colocalization. **(C,D**) Nuclei are labeled with DAPI (blue). Scale bars indicate 10µm.

## References

Aebi M. 2013. N-linked protein glycosylation in the ER. Biochim Biophys Acta - Mol Cell Res 1833:2430–2437. doi:10.1016/j.bbamcr.2013.04.001

Agrawal P, Fontanals-Cirera B, Sokolova E, Jacob S, Vaiana CA, Argibay D, Davalos V, McDermott M, Nayak S, Darvishian F, Castillo M, Ueberheide B, Osman I, Fenyö D, Mahal LK, Hernando E. 2017. A Systems Biology Approach Identifies FUT8 as a Driver of Melanoma Metastasis. Cancer Cell 31:804–819.e7. doi:10.1016/j.ccell.2017.05.007

Aoki K, Porterfield M, Lee SS, Dong B, Nguyen K, McGlamry KH, Tiemeyer M. 2008. The diversity of O-linked glycans expressed during Drosophila melanogaster development reflects stage-and tissue-specific requirements for cell signaling. J Biol Chem 283:30385–30400. doi:10.1074/jbc.M804925200

Aoki K, Tiemeyer M. 2010. The glycomics of glycan glucuronylation in drosophila melanogaster, 1st ed, Methods in Enzymology. Elsevier Inc. doi:10.1016/S0076-6879(10)80014-X

Aumiller JJ, Jarvis DL. 2002. Expression and functional characterization of a nucleotide sugar transporter from Drosophila melanogaster: relevance to protein glycosylation in insect cell expression systems. Protein Expr Purif 26:438–48.

Bach D-H, Park HJ, Lee SK. 2018. The Dual Role of Bone Morphogenetic Proteins in Cancer. Mol Ther oncolytics 8:1–13. doi:10.1016/j.omto.2017.10.002

Baldus SE, Zirbes TK, Hanisch F, Ph D, Kunze D, Shafizadeh ST, Nolden S, Mo SP, Karsten U, Ph D, Thiele J, Ho AH. 2000. Thomsen-Friedenreich Antigen Presents as a Prognostic Factor in Colorectal Carcinoma A Clinicopathologic Study of 264 Patients. Cancer 88:1536–1543.

Barr N, Taylor CR, Young T, Springer GF. 1989. Are pancarcinoma T and Tn differentiation antigens? Cancer 64:834–41.

Bennett EP, Chen YW, Schwientek T, Mandel U, Schjoldager KTBG, Cohen SM, Clausen H. 2010. Rescue of Drosophila Melanogaster l(2)35Aa lethality is only mediated by polypeptide GalNAc-transferase pgant35A, but not by the evolutionary conserved human ortholog GalNAc-transferase-T11. Glycoconj J 27:435–444. doi:10.1007/s10719-010-9290-5

Bennett EP, Mandel U, Clausen H, Gerken TA, Fritz TA, Tabak LA. 2012. Control of mucin-type O-glycosylation: a classification of the polypeptide GalNAc-transferase gene family. Glycobiology 22:736–56. doi:10.1093/glycob/cwr182

Bian B, Mongrain S, Cagnol S, Langlois M-J, Boulanger J, Bernatchez G, Carrier JC, Boudreau F, Rivard N. 2016. Cathepsin B promotes colorectal tumorigenesis, cell invasion, and metastasis. Mol Carcinog 55:671–87. doi:10.1002/mc.22312

Boersema PJ, Raijmakers R, Lemeer S, Mohammed S, Heck AJR. 2009. Multiplex peptide stable isotope dimethyl labeling for quantitative proteomics. Nat Protoc 4:484–494. doi:10.1038/nprot.2009.21

Boland CR, Montgomery CK, Kim YS. 1982. Alterations in human colonic mucin occurring with cellular differentiation and malignant transformation. Proc Natl Acad Sci U S A 79:2051–5.

Boskovski MT, Yuan S, Borbye Pedersen N, Knak Goth C, Makova S, Clausen H, Brueckner M, Khokha MF. 2015. The heterotaxy gene, GALNT11, glycosylates Notch to orchestrate cilia type and laterality. Anal Chem 25:368–379. doi:10.1016/j.cogdev.2010.08.003.Personal

Breloy I, Schwientek T, Althoff D, Holz M, Koppen T, Krupa A, Hanisch F-G. 2016. Functional Analysis of the Glucuronyltransferases GlcAT-P and GlcAT-S of Drosophila melanogaster: Distinct Activities towards the O-linked T-antigen. Biomolecules 6:8. doi:10.3390/biom6010008

Bruckner K, Kockel L, Duchek P, Luque CM, Rørth P, Perrimon N. 2004. The PDGF / VEGF Receptor Controls Blood Cell Survival in Drosophila 77 Avenue Louis Pasteur 7:73–84.

Campos-Ortega JA, Hartenstein V. 1997. The Embryonic Development of Drosophila melanogaster. Berlin, Heidelberg: Springer Berlin Heidelberg. doi:10.1007/978-3-662-22489-2

Cao B, Yang L, Rong W, Feng L, Han N, Zhang K, Cheng S, Wu J, Xiao T, Gao Y. 2015. Latent transforming growth factor-beta binding protein-1 in circulating plasma as a novel biomarker for early detection of hepatocellular carcinoma. Int J Clin Exp Pathol 8:16046–54.

Cao Y, Stosiek P, Springer GF, Karsten U. 1996. Thomsen-Friedenreich-related carbohydrate antigens in normal adult human tissues: a systematic and comparative study. Histochem Cell Biol 106:197–207.

Carrasco C, Gilhooly NS, Dillingham MS, Moreno-Herrero F. 2013. On the mechanism of recombination hotspot scanning during double-stranded DNA break resection. Proc Natl Acad Sci U S A 110:E2562–71. doi:10.1073/pnas.1303035110

Cawthorn TR, Moreno JC, Dharsee M, Tran-Thanh D, Ackloo S, Zhu PH, Sardana G, Chen J, Kupchak P, Jacks LM, Miller NA, Youngson BJ, Iakovlev V, Guidos CJ, Vallis KA, Evans KR, McCready D, Leong WL, Done SJ. 2012. Proteomic analyses reveal high expression of decorin and endoplasmin (HSP90B1) are associated with breast cancer metastasis and decreased survival. PLoS One 7:e30992. doi:10.1371/journal.pone.0030992

Chakravarthi S, Jessop CE, Willer M, Stirling CJ, Bulleid NJ. 2007. Intracellular catalysis of disulfide bond formation by the human sulfhydryl oxidase, QSOX1. Biochem J 404:403–411. doi:10.1042/BJ20061510

Chapel A, Kieffer-Jaquinod S, Sagné C, Verdon Q, Ivaldi C, Mellal M, Thirion J, Jadot M, Bruley C, Garin J, Gasnier B, Journet A. 2013. An Extended Proteome Map of the Lysosomal Membrane Reveals Novel Potential Transporters. Mol Cell Proteomics 12:1572–1588. doi:10.1074/mcp.M112.021980

Chia J, Goh G, Bard F. 2016. Short O-GalNAc glycans: regulation and role in tumor development and clinical perspectives. Biochim Biophys Acta - Gen Subj 1860:1623–1639. doi:10.1016/J.BBAGEN.2016.03.008

Chiu C-C, Lin C-Y, Lee L-Y, Chen Y-J, Lu Y-C, Wang H-M, Liao C-T, Chang JT-C, Cheng A-J. 2011. Molecular chaperones as a common set of proteins that regulate the invasion phenotype of head and neck cancer. Clin Cancer Res 17:4629–41. doi:10.1158/1078-0432.CCR-10-2107

Chiu C, Lin C, Lee L, Chen Y, Lu Y, Wang H. 2011. Molecular Chaperones as a Common Set of Proteins That Regulate the Invasion Phenotype of Head and Neck Cancer. Clin Cancer Res 17:1–14. doi:10.1158/1078-0432.CCR-10-2107

Christlet HT, Veluraja K. 2001. Database Analysis of O-Glycosylation Sites in Proteins. Biophys J 80:952–960. doi:10.1016/S0006-3495(01)76074-2

Cofre J, Abdelhay E. 2017. Cancer Is to Embryology as Mutation Is to Genetics□: Hypothesis of the Cancer as Embryological Phenomenon 2017. doi:10.1155/2017/3578090

Comer FI, Hart GW. 2000. O-glycosylation of nuclear and cytosolic proteins. Dynamic interplay between O-GlcNAc and O-phosphate. J Biol Chem 275:29179–29182. doi:10.1074/jbc.R000010200

Dalziel M, Whitehouse C, Mcfarlane I, Brockhausen I, Gschmeissner S, Schwientek T, Clausen H, Burchell JM, Taylor-papadimitriou J. 2001. The Relative Activities of the C2GnT1 and ST3Gal-I Glycosyltransferases Determine O-Glycan Structure and Expression of a Tumor-associated Epitope on MUC1 * 276:11007–11015. doi:10.1074/jbc.M006523200

Evans IR, Hu N, Skaer H, Wood W. 2010. Interdependence of macrophage migration and ventral nerve cord development in Drosophila embryos. Development 137:1625–1633. doi:10.1242/dev.046797

Fan X, Wang C, Song X, Liu H, Li X, Zhang Y. 2018. Elevated Cathepsin K potentiates metastasis of epithelial ovarian cancer. Histol Histopathol 33:673–680. doi:10.14670/HH-11-960

Ferguson K, Yadav A, Morey S, Abdullah J, Hrysenko G, Eng JY, Sajjad M, Koury S. 2014. Preclinical studies with JAA - F11 anti - Thomsen-Friedenreich monoclonal antibody for human breast cancer 10:385–399.

Fetchko M, Huang W, Li Y, Lai ZC. 2002. Drosophila Gp150 is required for early ommatidial development through modulation of Notch signaling. EMBO J 21:1074–1083. doi:10.1093/emboj/21.5.1074

Fu C, Zhao H, Wang Y, Cai H, Xiao Y, Zeng Y, Chen H. 2016. Tumor-associated antigens: Tn antigen, sTn antigen, and T antigen. Hla 88:275–286. doi:10.1111/tan.12900

Fuwa TJ, Kinoshita T, Nishida H, Nishihara S. 2015. Reduction of T antigen causes loss of hematopoietic progenitors in Drosophila through the inhibition of filopodial extensions from the hematopoietic niche. Dev Biol 401:206–219. doi:10.1016/j.ydbio.2015.03.003

Garcia A, Kandel JJ. 2012. Notch: a key regulator of tumor angiogenesis and metastasis. Histol Histopathol 27:151–6. doi:10.14670/HH-27.151

Gohrig A, Detjen KM, Hilfenhaus G, Korner JL, Welzel M, Arsenic R, Schmuck R, Bahra M, Wu JY, Wiedenmann B, Fischer C. 2014. Axon Guidance Factor SLIT2 Inhibits Neural Invasion and Metastasis in Pancreatic Cancer. Cancer Res 74:1529–1540. doi:10.1158/0008-5472.CAN-13-1012

Gonias SL, Karimi-Mostowfi N, Murray SS, Mantuano E, Gilder AS. 2017. Expression of LDL receptor-related proteins (LRPs) in common solid malignancies correlates with patient survival. PLoS One 12:e0186649. doi:10.1371/journal.pone.0186649

Goth CK, Vakhrushev SY, Joshi HJ, Clausen H, Schjoldager KT. 2018. Fine-Tuning Limited Proteolysis: A Major Role for Regulated Site-Specific O-Glycosylation. Trends Biochem Sci 43:269–284. doi:10.1016/j.tibs.2018.02.005

Guruharsha KG, Rual JF, Zhai B, Mintseris J, Vaidya P, Vaidya N, Beekman C, Wong C, Rhee DY, Cenaj O, McKillip E, Shah S, Stapleton M, Wan KH, Yu C, Parsa B, Carlson JW, Chen X, Kapadia B, Vijayraghavan K, Gygi SP, Celniker SE, Obar RA, Artavanis-Tsakonas S. 2011. A protein complex network of Drosophila melanogaster. Cell 147:690–703. doi:10.1016/j.cell.2011.08.047

Gyoergy A, Roblek M, Ratheesh A, Valoskova K, Belyaeva V, Wachner S, Matsubayashi Y, Sánchez-Sánchez BJ, Stramer B, Siekhaus DE. 2018. Tools Allowing Independent Visualization and Genetic Manipulation of Drosophila melanogaster Macrophages and Surrounding Tissues. G3 (Bethesda) 8:845–857. doi:10.1534/g3.117.300452

Hamaratoglu F, Affolter M, Pyrowolakis G. 2014. Dpp/BMP signaling in flies: From molecules to biology. Semin Cell Dev Biol 32:128–136. doi:10.1016/j.semcdb.2014.04.036

Hart GW, Slawson C, Ramirez-Correa G, Lagerlof O. 2011. Cross talk between O-GlcNAcylation and phosphorylation: roles in signaling, transcription, and chronic disease. Annu Rev Biochem 80:825–58. doi:10.1146/annurev-biochem-060608-102511

Hassan H, Reis CA, Bennett EP, Mirgorodskaya E, Roepstorff P, Hollingsworth MA, Burchell J, Taylor-Papadimitriou J, Clausen H. 2000. The lectin domain of UDP-N-acetyl-D-galactosamine: polypeptide N-acetylgalactosaminyltransferase-T4 directs its glycopeptide specificities. J Biol Chem 275:38197–205. doi:10.1074/jbc.M005783200

Häuselmann I, Borsig L. 2014. Altered tumor-cell glycosylation promotes metastasis. Front Oncol 4:1–15. doi:10.3389/fonc.2014.00028

Heimburg J, Yan J, Morey S, Glinskii O V., Huxley VH, Wild L, Klick R, Roy R, Glinsky V V., Rittenhouse-Olson K. 2006. Inhibition of Spontaneous Breast Cancer Metastasis by Anti—Thomsen-Friedenreich Antigen Monoclonal Antibody JAA-F11. Neoplasia 8:939–948. doi:10.1593/neo.06493

Hodar C, Zuñiga A, Pulgar R, Travisany D, Chacon C, Pino M, Maass A, Cambiazo V. 2014. Comparative gene expression analysis of Dtg, a novel target gene of Dpp signaling pathway in the early Drosophila melanogaster embryo. Gene 535:210–217. doi:10.1016/j.gene.2013.11.032

Hofman K, Stoffel W. 1993. TMbase-A database of membrane spanning proteins segments. BiolChem 374:166.

Howard DR, Taylor CR. 1980. An antitumor antibody in normal human serum: reaction of anti-T with breast carcinoma cells. Oncology 37:142–8. doi:10.1159/000225423

Hu X, Wang Q, Tang M, Barthel F, Amin S, Yoshihara K, Lang FM, Martinez-ledesma E, Lee SH, Zheng S, Verhaak RGW. 2018. TumorFusions□: an integrative resource for cancer-associated transcript fusions. Nucleic Acids Res 46:1144–1149. doi:10.1093/nar/gkx1018

Huang X, Jin M, Chen Y-X, Wang J, Zhai K, Chang Y, Yuan Q, Yao K-T, Ji G. 2016. ERP44 inhibits human lung cancer cell migration mainly via IP3R2. Aging (Albany NY) 8:1276–86. doi:10.18632/aging.100984

Hung J-S, Huang J, Lin Y-C, Huang M-J, Lee P-H, Lai H-S, Liang J-T, Huang M-C. 2014. C1GALT1 overexpression promotes the invasive behavior of colon cancer cells through modifying O-glycosylation of FGFR2. Oncotarget 5:2096–106. doi:10.18632/oncotarget.1815

Itoh K, Akimoto Y, Fuwa TJ, Sato C, Komatsu A. 2016. Mucin-type core 1 glycans regulate the localization of neuromuscular junctions and establishment of muscle cell architecture in Drosophila. Dev Biol 412:114–127. doi:10.1016/j.ydbio.2016.01.032

Itoh K, Akimoto Y, Kondo S, Ichimiya T, Aoki K. 2018. Glucuronylated core 1 glycans are required for precise localization of neuromuscular junctions and normal formation of basement membranes on Drosophila muscles. Dev Biol 436:108–124. doi:10.1016/j.ydbio.2018.02.017

Ju T, Cummings RD. 2005. Protein glycosylation: Chaperone mutation in Tn syndrome. Nature 437:1252–1252. doi:10.1038/4371252a

Katchman BA, Antwi K, Hostetter G, Demeure MJ, Watanabe A, Decker GA, Miller LJ, Von Hoff DD, Lake DF. 2011. Quiescin sulfhydryl oxidase 1 promotes invasion of pancreatic tumor cells mediated by matrix metalloproteinases. Mol Cancer Res 9:1621–31. doi:10.1158/1541-7786.MCR-11-0018

Katchman BA, Ocal IT, Cunliffe HE, Chang Y, Hostetter G, Watanabe A, Lobello J, Lake DF. 2013. Expression of quiescin sulfhydryl oxidase 1 is associated with a highly invasive phenotype and correlates with a poor prognosis in Luminal B breast cancer Expression of quiescin sulfhydryl oxidase 1 is associated with a highly invasive phenotype and corre. Breast Cancer Res 15:R28. doi:10.1186/bcr3407

Kellokumpu S, Sormunen R, Kellokumpu I. 2002. Abnormal glycosylation and altered Golgi structure in colorectal cancer: dependence on intra-Golgi pH. FEBS Lett 516:217–24.

Kim B-T, Tsuchida K, Lincecum J, Kitagawa H, Bernfield M, Sugahara K. 2003. Identification and characterization of three Drosophila melanogaster glucuronyltransferases responsible for the synthesis of the conserved glycosaminoglycan-protein linkage region of proteoglycans. Two novel homologs exhibit broad specificity toward oligosaccharides from proteoglycans, glycoproteins, and glycosphingolipids. J Biol Chem 278:9116–24. doi:10.1074/jbc.M209344200

Kornfeld R, Kornfeld S. 1985. Assembly of asparagine-linked oligosaccharides. Annu Rev Biochem 54:631–64. doi:10.1146/annurev.bi.54.070185.003215

Kubota T, Shiba T, Sugioka S, Furukawa S, Sawaki H, Kato R, Wakatsuki S, Narimatsu H. 2006. Structural basis of carbohydrate transfer activity by human UDP-GalNAc: polypeptide alpha-N-acetylgalactosaminyltransferase (pp-GalNAc-T10). J Mol Biol 359:708–27. doi:10.1016/j.jmb.2006.03.061

Lake DF, Faigel DO. 2014. The Emerging Role of QSOX1 in Cancer. Antioxid Redox Signal 21:485–496. doi:10.1089/ars.2013.5572

Lehmann R, Tautz D. 1994. In Situ Hybridization to RNA. Methods Cell Biol 44:575–598. doi:10.1016/S0091-679X(08)60933-4

Letsou A, Arora K, Wrana JL, Simin K, Twombly V, Jamal J, Staehling-Hampton K, Hoffmann FM, Gelbart WM, Massagué J, O’Connor MB. 1995. Drosophila Dpp signaling is mediated by the punt gene product: A dual ligand-binding type II receptor of the TGFβ receptor family. Cell 80:899–908. doi:10.1016/0092-8674(95)90293-7

Levery SB, Steentoft C, Halim A, Narimatsu Y, Clausen H, Vakhrushev SY. 2015. Advances in mass spectrometry driven O-glycoproteomics. Biochim Biophys Acta - Gen Subj 1850:33–42. doi:10.1016/j.bbagen.2014.09.026

Li S, Liu P, Xi L, Jiang X, Wu M, Deng D, Wei J, Zhu T, Zhou L, Wang S, Xu G, Meng L, Zhou J, Ma D. 2008. Expression of TMEM87B interacting with the human papillomavirus type 18 E6 oncogene in the Hela cDNA library by a yeast two-hybrid system. Oncol Rep 20:421–7.

Li Y. 2003. Scabrous and Gp150 are endosomal proteins that regulate Notch activity. Development 130:2819–2827. doi:10.1242/dev.00495

Limas C, Lange P. 1986. T-antigen in normal and neoplastic urothelium. Cancer 58:1236–45.

Lin YR, Reddy BVVG, Irvine KD. 2008. Requirement for a core 1 galactosyltransferase in the Drosophila nervous system. Dev Dyn 237:3703–3714. doi:10.1002/dvdy.21775

Linton KM, Hey Y, Saunders E, Jeziorska M, Denton J, Wilson CL, Swindell R, Dibben S, Miller CJ, Pepper SD, Radford JA, Freemont AJ. 2008. Acquisition of biologically relevant gene expression data by Affymetrix microarray analysis of archival formalin-fixed paraffin-embedded tumours. Br J Cancer 98:1403–14. doi:10.1038/sj.bjc.6604316

MacLean GD, Longenecker BM. 1991. Clinical significance of the Thomsen-Friedenreich antigen. Semin Cancer Biol 2:433–9.

Mao F, Holmlund C, Faraz M, Wang W, Bergenheim T, Kvarnbrink S, Johansson M, Henriksson R, Hedman H. 2018. Lrig1 is a haploinsufficient tumor suppressor gene in malignant glioma. Oncogenesis 7:1–12. doi:10.1038/s41389-017-0012-8

Marshall R. 1972. Glycoproteins. Annu Rev Biochem 41:673–702.

Matos LL, Suarez ER, Theodoro TR, Trufelli DC, Melo CM, Garcia LF, Oliveira OCG, Matos MGL, Kanda JL, Nader HB, Martins JRM, Pinhal MAS. 2015. The Profile of Heparanase Expression Distinguishes Differentiated Thyroid Carcinoma from Benign Neoplasms. PLoS One 10:e0141139. doi:10.1371/journal.pone.0141139

Meerbrey KL, Hu G, Kessler JD, Roarty K, Li MZ, Fang JE, Herschkowitz JI, Burrows AE, Ciccia A, Sun T, Schmitt EM, Bernardi RJ, Fu X, Bland CS, Cooper TA, Schiff R, Rosen JM, Westbrook TF, Elledge SJ. 2011. The pINDUCER lentiviral toolkit for inducible RNA interference in vitro and in vivo. Proc Natl Acad Sci 108:3665–3670. doi:10.1073/pnas.1019736108

Molin K, Fredman P, Svennerholm L. 1986. Binding specificities of the lectins PNA, WGA and UEA I to polyvinylchloride-adsorbed glycosphingolipids. FEBS Lett 205:51–55. doi:10.1016/0014-5793(86)80864-X

Müller R, Hülsmeier AJ, Altmann F, Ten Hagen K, Tiemeyer M, Hennet T. 2005. Characterization of mucin-type core-1 β1-3 galactosyltransferase homologous enzymes in Drosophila melanogaster. FEBS J 272:4295–4305. doi:10.1111/j.1742-4658.2005.04838.x

Natchiar SK, Suguna K, Surolia A, Vijayan M. 2007. Peanut agglutinin, a lectin with an unusual quaternary structure and interesting ligand binding properties. Crystallogr Rev 13:3–28. doi:10.1080/08893110701382087

Ninov N, Menezes-Cabral S, Prat-Rojo C, Manjón C, Weiss A, Pyrowolakis G, Affolter M, Martín-Blanco E. 2010. Dpp signaling directs cell motility and invasiveness during epithelial morphogenesis. Curr Biol 20:513–20. doi:10.1016/j.cub.2010.01.063

Ntziachristos P, Lim JS, Sage J, Aifantis I. 2014. From Fly Wings to Targeted Cancer Therapies: A Centennial for Notch Signaling. Cancer Cell 25:318–334. doi:10.1016/j.ccr.2014.02.018

Ohtsubo K, Marth JD. 2006. Glycosylation in cellular mechanisms of health and disease. Cell 126:855–67. doi:10.1016/j.cell.2006.08.019

Omasits U, Ahrens CH, Müller S, Wollscheid B. 2014. Protter: Interactive protein feature visualization and integration with experimental proteomic data. Bioinformatics 30:884–886. doi:10.1093/bioinformatics/btt607

Orntoft TF, Mors NP, Eriksen G, Jacobsen NO, Poulsen HS. 1985. Comparative immunoperoxidase demonstration of T-antigens in human colorectal carcinomas and morphologically abnormal mucosa. Cancer Res 45:447–52.

Owens P, Pickup MW, Novitskiy S V, Giltnane JM, Gorska AE, Hopkins CR, Hong CC, Moses HL. 2015. Inhibition of BMP signaling suppresses metastasis in mammary cancer. Oncogene 34:2437–49. doi:10.1038/onc.2014.189

Pacquelet A, Røth P. 1999. Regulatory mechanisms required for DE-cadherin function in cell migration and other types of adhesion. J Cell Biol 170:803–812. doi:10.1083/jcb.200506131

Palmieri M, Impey S, Kang H, di Ronza A, Pelz C, Sardiello M, Ballabio A. 2011. Characterization of the CLEAR network reveals an integrated control of cellular clearance pathways. Hum Mol Genet 20:3852–3866. doi:10.1093/hmg/ddr306

Pickup MW, Hover LD, Guo Y, Gorska AE, Chytil A, Novitskiy S V, Moses HL, Owens P. 2015. Deletion of the BMP receptor BMPR1a impairs mammary tumor formation and metastasis. Oncotarget 6:22890–904. doi:10.18632/oncotarget.4413

Pierce GB. 1974. Neoplasms, differentiations and mutations. Am J Pathol 77:103–118.

Piller V, Piller F, Cartron J. 1990. Comparison of the carbohydrate-binding specificities of seven N-acetyl-D-galactosamine-recognizing lectins. Eur J Biochem 191:461–466. doi:10.1111/j.1432-1033.1990.tb19144.x

Quistgaard EM, Löw C, Guettou F, Nordlund P. 2016. Understanding transport by the major facilitator superfamily (MFS): structures pave the way. Nat Rev Mol Cell Biol 17:123–132. doi:10.1038/nrm.2015.25

Ratheesh A, Biebl J, Vesela J, Smutny M, Papusheva E, Krens SFG, Kaufmann W, Gyoergy A, Casano AM, Siekhaus DE. 2018. Drosophila TNF Modulates Tissue Tension in the Embryo to Facilitate Macrophage Invasive Migration. Dev Cell 45:331–346.e7. doi:10.1016/j.devcel.2018.04.002

Reddy VS, Shlykov MA, Castillo R, Sun EI, Saier MH. 2012. The major facilitator superfamily (MFS) revisited. FEBS J 279:2022–2035. doi:10.1111/j.1742-4658.2012.08588.x

Revoredo L, Wang S, Bennett EP, Clausen H, Moremen KW, Jarvis DL, Ten Hagen KG, Tabak LA, Gerken TA. 2016. Mucin-type o-glycosylation is controlled by short-And long-range glycopeptide substrate recognition that varies among members of the polypeptide GalNAc transferase family. Glycobiology 26:360–376. doi:10.1093/glycob/cwv108

Riedel F, Gillingham AK, Rosa-Ferreira C, Galindo A, Munro S. 2016. An antibody toolkit for the study of membrane traffic in *Drosophila melanogaster*. Biol Open 5:987–992. doi:10.1242/bio.018937

Sahlgren C, Gustafsson M V, Jin S, Poellinger L, Lendahl U. 2008. Notch signaling mediates hypoxia-induced tumor cell migration and invasion. Proc Natl Acad Sci U S A 105:6392–7. doi:10.1073/pnas.0802047105

Schindlbeck C, Jeschke U, Schulze S, Karsten U, Janni W, Rack B, Sommer H, Friese K. 2005. Characterisation of disseminated tumor cells in the bone marrow of breast cancer patients by the Thomsen–Friedenreich tumor antigen. Histochem Cell Biol 123:631–637. doi:10.1007/s00418-005-0781-6

Schjoldager KT-BG, Vakhrushev SY, Kong Y, Steentoft C, Nudelman AS, Pedersen NB, Wandall HH, Mandel U, Bennett EP, Levery SB, Clausen H. 2012. Probing isoform-specific functions of polypeptide GalNAc-transferases using zinc finger nuclease glycoengineered SimpleCells. Proc Natl Acad Sci 109:9893–9898. doi:10.1073/pnas.1203563109

Schjoldager KT, Joshi HJ, Kong Y, Goth CK, King SL, Wandall HH, Bennett EP, Vakhrushev SY, Clausen H. 2015. Deconstruction of O-glycosylation--GalNAc-T isoforms direct distinct subsets of the O-glycoproteome. EMBO Rep 16:1713–1722. doi:10.15252/embr.201540796

Schwientek T, Bennett EP, Flores C, Thacker J, Hollmann M, Reis CA, Behrens J, Mandel U, Keck B, Mireille A, Schäfer, Haselmann K, Zubarev R, Roepstorff P, Burchell JM, Taylor-Papadimitriou J, Hollingsworth MA, Clausena H. 2002. Functional conservation of subfamilies of putative UDP-N-acetylgalactosamine:Polypeptide N-acetylgalactosaminyltransferases in Drosophila, Caenorhabditis elegans, and mammals. One subfamily composed of l(2)35Aa is essential in Drosophila. J Biol Chem 277:22623–22638. doi:10.1074/jbc.M202684200

Schwientek T, Mandel U, Roth U, Müller S, Hanisch FG. 2007. A serial lectin approach to the mucin-type O-glycoproteome of Drosophila melanogaster S2 cells. Proteomics 7:3264–3277. doi:10.1002/pmic.200600793

Segawa H, Kawakita M, Ishida N. 2002. Human and Drosophila UDP-galactose transporters transport UDP-N-acetylgalactosamine in addition to UDP-galactose. Eur J Biochem 269:128–38.

Senanayake U, Das S, Vesely P, Alzoughbi W, Frohlich LF, Chowdhury P, Leuschner I, Hoefler G, Guertl B. 2012. miR-192, miR-194, miR-215, miR-200c and miR-141 are downregulated and their common target ACVR2B is strongly expressed in renal childhood neoplasms. Carcinogenesis 33:1014–1021. doi:10.1093/carcin/bgs126

Sheu JJ-C, Lee C-C, Hua C-H, Li C-I, Lai M-T, Lee S-C, Cheng J, Chen C-M, Chan C, Chao SC-C, Chen J-Y, Chang J-Y, Lee C-H. 2014. LRIG1 modulates aggressiveness of head and neck cancers by regulating EGFR-MAPK-SPHK1 signaling and extracellular matrix remodeling. Oncogene 33:1375–1384. doi:10.1038/onc.2013.98

Siekhaus D, Haesemeyer M, Moffitt O, Lehmann R. 2010. RhoL controls invasion and Rap1 localization during immune cell transmigration in Drosophila. Nat Cell Biol 12:605–610. doi:10.1038/ncb2063

Sonoshita M, Aoki M, Fuwa H, Aoki K, Hosogi H, Sakai Y, Hashida H, Takabayashi A, Sasaki M, Robine S, Itoh K, Yoshioka K, Kakizaki F, Kitamura T, Oshima M, Taketo MM. 2011. Suppression of colon cancer metastasis by Aes through inhibition of Notch signaling. Cancer Cell 19:125–37. doi:10.1016/j.ccr.2010.11.008

Springer GF. 1997. Immunoreactive T and Tn epitopes in cancer diagnosis, prognosis, and immunotherapy 594–602.

Springer GF. 1989. Tn epitope (N-acetyl-D-galactosamine alpha-O-serine/threonine) density in primary breast carcinoma: a functional predictor of aggressiveness. Mol Immunol 26:1–5.

Springer GF. 1984. T and Tn, General Carcinoma Autoantigens. Science (80-) 224:1198–1206.

Springer GF, Desai PR, Banatwala I. 1975. Blood group MN antigens and precursors in normal and malignant human breast glandular tissue. J Natl Cancer Inst 54:335–9.

Stanley P, Schachter H, Taniguchi N. 2009. N-Glycans, Essentials of Glycobiology. Cold Spring Harbor Laboratory Press.

Steentoft C, Vakhrushev SY, Joshi HJ, Kong Y, Vester-Christensen MB, Schjoldager KTBG, Lavrsen K, Dabelsteen S, Pedersen NB, Marcos-Silva L, Gupta R, Paul Bennett E, Mandel U, Brunak S, Wandall HH, Levery SB, Clausen H. 2013. Precision mapping of the human O-GalNAc glycoproteome through SimpleCell technology. EMBO J 32:1478–1488. doi:10.1038/emboj.2013.79

Steentoft C, Vakhrushev SY, Vester-Christensen MB, Schjoldager KT-BG, Kong Y, Bennett EP, Mandel U, Wandall H, Levery SB, Clausen H. 2011. Mining the O-glycoproteome using zinc-finger nuclease–glycoengineered SimpleCell lines. Nat Methods 8:977–982. doi:10.1038/nmeth.1731

Stojadinovic A, Hooke JA, Shriver CD, Nissan A, Kovatich AJ, Kao T-C, Ponniah S, Peoples GE, Moroni M. 2007. HYOU1 / Orp150 expression in breast cancer. Med Sci Monit 13:231–239.

Summers JL, Coon JS, Ward RM, Falor WH, Miller AW, Weinstein RS. 1983. Prognosis in carcinoma of the urinary bladder based upon tissue blood group abh and Thomsen-Friedenreich antigen status and karyotype of the initial tumor. Cancer Res 43:934–9.

Tarp MA, Clausen H. 2008. Mucin-type O-glycosylation and its potential use in drug and vaccine development. Biochim Biophys Acta 1780:546–63. doi:10.1016/j.bbagen.2007.09.010

Ten Hagen KG, Tran DT, Gerken TA, Stein DS, Zhang Z. 2003. Functional characterization and expression analysis of members of the UDP-GalNAc:polypeptide N-acetylgalactosaminyltransferase family from Drosophila melanogaster. J Biol Chem 278:35039–35048. doi:10.1074/jbc.M303836200

Tepass U, Eileen G, F D, Haag A, Omatyar L, Trk T. 1996. shotgun encodes Drosophila E-cadherin and is preferentially, required during cell rearrangement in the neurectoderm and other morphogenetically active epithelia 672–685.

Tian E, Ten Hagen KG. 2009. Recent insights into the biological roles of mucin-type O-glycosylation. Glycoconj J 26:325–34. doi:10.1007/s10719-008-9162-4

Tian E, Ten Hagen KG. 2006. Expression of the UDP-GalNAc: Polypeptide N-acetylgalactosaminyltransferase family is spatially and temporally regulated during Drosophila development. Glycobiology 16:83–95. doi:10.1093/glycob/cwj051

Tien AC, Rajan A, Schulze KL, Hyung DR, Acar M, Steller H, Bellen HJ. 2008. Ero1L, a thiol oxidase, is required for Notch signaling through cysteine bridge formation of the Lin12-Notch repeats in Drosophila melanogaster. J Cell Biol 182:1113–1125. doi:10.1083/jcb.200805001

Tomancak P, Beaton A, Weiszmann R, Kwan E, Shu S, Lewis SE, Richards S, Ashburner M, Hartenstein V, Celniker SE, Rubin GM. 2002. Systematic determination of patterns of gene expression during Drosophila embryogenesis. Genome Biol 3:RESEARCH0088. doi:10.1186/gb-2002-3-12-research0088

Tomancak P, Berman BP, Beaton A, Weiszmann R, Kwan E, Hartenstein V, Celniker SE, Rubin GM. 2007. Global analysis of patterns of gene expression during Drosophila embryogenesis 8:1–24. doi:10.1186/gb-2007-8-7-r145

Tran DT, Zhang L, Zhang Y, Tian E, Earl LA, Ten Hagen KG. 2012. Multiple members of the UDP-GalNAc: Polypeptide N-acetylgalactosaminyltransferase family are essential for viability in Drosophila. J Biol Chem 287:5243–5252. doi:10.1074/jbc.M111.306159

Varki A. 2017. Biological roles of glycans. Glycobiology 27:3–49. doi:10.1093/glycob/cww086

Verheyen EM, Cooley L. 1994. Profilin mutations disrupt multiple actin-dependent processes during Drosophila development. Development 120:717–28.

Vizcaino JA, Deutch EW, Wang R, Csordas A, Raisinger F, Rios D, Dianes JA, Sun Z, Farrah T, Bandeira N, Binz P-A, Xenarios I, Eisenacher M, Mayer G, Gatto L, Campos A, Chalkley RJ, Kraus H-J, Albar PJ, Martinez-Bartolome S, Apweiler R, Omenn GS, Martens L, Jones AR, Hermjakob H. 2016. ProteomeXchange provides globally co-ordinated proteomics data submission and dissemination Juan 10:1939–1947. doi:10.1038/nprot.2015.121.Human

Walls G V, Stevenson M, Lines KE, Newey PJ, Reed AAC, Bowl MR, Jeyabalan J, Harding B, Bradley KJ, Manek S, Chen J, Wang P, Williams BO, Teh BT, Thakker R V. 2017. Mice deleted for cell division cycle 73 gene develop parathyroid and uterine tumours: model for the hyperparathyroidism-jaw tumour syndrome. Oncogene 36:4025–4036. doi:10.1038/onc.2017.43

Webb DJ, Nguyen DH, Sankovic M, Gonias SL. 1999. The very low density lipoprotein receptor regulates urokinase receptor catabolism and breast cancer cell motility in vitro. J Biol Chem 274:7412–20.

Yokdang N, Hatakeyama J, Wald JH, Simion C, Tellez JD, Chang DZ, Swamynathan MM, Chen M, Murphy WJ, Carraway Iii KL, Sweeney C. 2016. LRIG1 opposes epithelial-to-mesenchymal transition and inhibits invasion of basal-like breast cancer cells. Oncogene 35:2932–47. doi:10.1038/onc.2015.345

Yoshida H, Fuwa TJ, Arima M, Hamamoto H, Sasaki N, Ichimiya T, Osawa KI, Ueda R, Nishihara S. 2008. Identification of the Drosophila core 1 β1,3-galactosyltransferase gene that synthesizes T antigen in the embryonic central nervous system and hemocytes. Glycobiology 18:1094–1104. doi:10.1093/glycob/cwn094

Yu LG, Andrews N, Zhao Q, McKean D, Williams JF, Connor LJ, Gerasimenko O V., Hilkens J, Hirabayashi J, Kasai K, Rhodes JM. 2007. Galectin-3 interaction with Thomsen-Friedenreich disaccharide on cancer-associated MUC1 causes increased cancer cell endothelial adhesion. J Biol Chem 282:773–781. doi:10.1074/jbc.M606862200

Zhang L, Tran DT, Ten Hagen KG. 2010. An O-glycosyltransferase promotes cell adhesion during development by influencing secretion of an extracellular matrix integrin ligand. J Biol Chem 285:19491–19501. doi:10.1074/jbc.M109.098145

Zhang Y, Ran Y, Xiong Y, Zhong Z-B, Wang Z-H, Fan X-L, Ye Q-F. 2016. Effects of TMEM9 gene on cell progression in hepatocellular carcinoma by RNA interference. Oncol Rep 36:299–305. doi:10.3892/or.2016.4821

Zhou Y, Liao Q, Li X, Wang H, Wei F, Chen J, Yang J. 2016. HYOU1, Regulated by LPLUNC1, Is Up-Regulated in Nasopharyngeal Carcinoma and Associated with Poor Prognosis. J Cancer 7:367–367. doi:10.7150/jca.13695

Zhou Y, Zhu Y, Fan X, Zhang C, Wang Y, Zhang L, Zhang H, Wen T, Zhang K, Huo X, Jiang X, Bu Y, Zhang Y. 2017. NID1, a new regulator of EMT required for metastasis and chemoresistance of ovarian cancer cells. Oncotarget 8:33110–33121. doi:10.18632/oncotarget.16145

